# Hidden partners: Using cross-docking calculations to predict binding sites for proteins with multiple interactions

**DOI:** 10.1101/244913

**Authors:** Nathalie Lagarde, Alessandra Carbone, Sophie Sacquin-Mora

## Abstract

Protein-protein interactions control a large range of biological processes and their identification is essential to understand the underlying biological mechanisms. To complement experimental approaches, *in silico* methods are available to investigate protein-protein interactions. Cross-docking methods, in particular, can be used to predict protein binding sites. However, proteins can interact with numerous partners and can present multiple binding sites on their surface, which may alter the binding site prediction quality. We evaluate the binding site predictions obtained using complete cross-docking simulations of 358 proteins with two different scoring schemes accounting for multiple binding sites. Despite overall good binding site prediction performances, 68 cases were still associated with very low prediction quality, presenting individual area under the specificity-sensitivity ROC curve (AUC) values below the random AUC threshold of 0.5, since cross-docking calculations can lead to the identification of alternate protein binding sites (that are different from the reference experimental sites). For the large majority of these proteins, we show that the predicted alternate binding sites correspond to interaction sites with hidden partners, i.e. partners not included in the original cross-docking dataset. Among those new partners, we find proteins, but also nucleic acid molecules. Finally, for proteins with multiple binding sites on their surface, we investigated the structural determinants associated with the binding sites the most targeted by the docking partners.

**Abbreviations:** ANOVA: ANalysis Of Variance; AUC: Area Under the Curve; Best Interface: BI; CAPRI: Critical Assessment of Prediction of Interactions; CC-D: Complete Cross-Docking; DNA: DesoxyriboNucleic Acid; FDR: False Discovery Rate; FRI_res(type)_: Fraction of each Residue type in the Interface; FP: False Positives; GI: Global Interface; HCMD: Help Cure Muscular Dystrophy; JET: Joint Evolutionary Tree; MAXDo: Molecular Association via Cross Docking; NAI: Nucleic Acid Interface; NPV: Negative Predicted Value; PDB: Protein Data Bank; PIP: Protein Interface Propensity; PiQSi: Protein Quaternary Structure investigation; PPIs: Protein-Protein Interactions; PPV: Positive Predicted Value; Prec.: Precision; PrimI: Primary Interface; RNA: RiboNucleic Acid; ROC: Receiver Operating Characteristic; SecI: Secondary Interface; Sen.: Sensitivity; Spe.: Specificity; TN: True Negatives; TP: True Positives; WCG: World Community Grid.

## Introduction

Proteins play a fundamental role in many biological process (metabolism, information processing, transport, structural organization), through physical interactions with other proteins and molecules such as metabolites, lipids and nucleic acids ^1^. In particular, protein-protein interactions (PPIs) control the assembly of proteins in large edifices forming complex molecular machines ^2^.

The study of PPIs permits to decipher the protein network constituting the interactome of an organism, to understand the *molecular sociology of the cell* ^3^ and to explain their role in biological systems ^4–8^. PPIs detection can be realized using numerous experimental approaches ^9^, including *in vitro* techniques such as tandem affinity purification and *in vivo* methods like the yeast two-hybrid. However, these experimental methods are associated with several limitations, such as cost, time, a low interaction coverage, and biases toward certain protein types and cellular localizations, that generate a significant number of false positives and negatives ^10^. On the other hand, *in silico* methods have been developed and constitute complementary approaches to experimental techniques ^11–13^. Molecular modeling can notably be used to identify protein interactions, with the advantage of providing structural models for the corresponding complexes and insights into the physical principles behind the complex formation. Docking methods, which were originally developed to predict the structure of a complex starting from the structures of two proteins known to interact ^14^, can be diverted for the prediction of protein interfaces. In this perspective, the collection of docking poses will be used to derive a consensus of predicted interface residues ^15^ both in single docking studies ^16–20^, in which the docking poses result from the docking of two protein partners already known to interact, and in complete cross-docking (CC-D) studies ^21–24^, which involve performing docking calculations on all possible protein pairs within a given dataset. Several benchmarking databases are available to evaluate protein-protein docking protocols, such as the Docking benchmark 5.0 ^25^, DOCKGROUND ^26^, or 2P2IDB ^27^. The Docking benchmark 5.0 ^25^ includes 230 protein-protein complexes and provides both bound and unbound structures for the majority of the protein partners. It is considered as the gold standard for protein-protein docking evaluation, notably thanks to the careful protein annotations (about the functional and docking difficulty categories). One should note that our docking algorithm was evaluated in a previous study on the 168 proteins of the Docking benchmark 2.0 ^28^. DOCKGROUND ^26^ is a large protein-protein interactions database, with 396 co-crystallized protein-protein complexes and the corresponding X-ray unbound structures for both proteins. 2P2Idb ^27^ is a smaller database dedicated to orthosteric modulation of protein-protein interactions. Its specificity is to provide all interactions for which both the protein-protein and protein-inhibitor have been structurally resolved, resulting in a database of 31 protein-protein complexes, 619 protein-ligand complexes and 553 PPI inhibitors. Earlier studies ^21, 23, 24^, using such benchmarking databases, showed that the cross-docking method could be used to accurately predict protein binding sites. However, proteins are able to interact with different partners, forming an intricate network and thus complicating PPI analysis ^29, 30^. In their work on over 35 000 protein complex structures from the PDB, Zhao et al. ^31^ found that around 40% of the proteins presented multiple interfaces, an estimation which concurs with the work of Kim et al. ^32^, who estimated that 40% of protein domains can bind via multiple orientation.

Among them, proteins presenting more than 5 partners are defined as *hub* (in reference to the protein interaction networks) or *social partners*. These hub proteins can be classified according to the number of binding sites exhibited on their surface into singlish interface hubs (one or two *multibinding protein interface* ^33^) or multi-interface hubs (larger number of binding sites) ^34^. To our knowledge the impact on cross-docking binding site predictions of the existence of multiple binding sites on the protein surface has never been systematically addressed. However, previous studies using a different approach based on sequence and structure analysis of a single protein, tackled the problem of predicting multiple binding sites for proteins ^35, 36^.

In the cross-docking dataset of 358 proteins used in this work, even if the majority of proteins were associated with only one experimental partner, some multi-interface proteins presenting up to 5 experimental partners were also available. The first aim of our study was to evaluate both the binding site predictions obtained using cross-docking simulations with our dataset and the ability of the cross-docking method to detect multiple binding sites on protein surfaces. Therefore, we compared the efficiency of two scoring schemes accounting for multiple binding sites.

In a second step, we analyzed the cases where cross-docking calculations lead to the identification of alternate protein binding sites that were different from the reference experimental sites based on the interaction partners present in the protein dataset. For the large majority of these proteins, the predicted alternate binding sites were shown to correspond to interaction sites with other partners not included in the original cross-docking dataset. Finally, we tried to understand why some interfaces were better predicted than others using our CC-D results, and why, in some cases, protein-protein docking results could lead to the identification of binding sites for nucleic acids.

## Materials and Methods

### Cross-docking calculations

#### Protein dataset

The HCMD2 dataset comprises 2246 non-redundant proteins. Among them around 400 are known to be involved in neuromuscular diseases (around 200 experimentally-determined structures and around 200 predicted structural models) and the 1800 remaining proteins, with a role in muscular dystrophy that is yet unknown, are human proteins, notably involved in the pathways monitoring essential cardiac or cerebral mechanisms. The HCMD dataset was constructed with the aim to study the potential interactions of the 400 proteins involved in neuromuscular diseases within the human body, to provide new insights on their molecular mechanism and to help biomedical researchers to develop therapies for neuromuscular diseases (for more information see Section 5 “Proteins list that will be analyzed on phase 2 of the HCMD2 project” in http://www.ihes.fr/~carbone/HCMDproject.htm). Starting from the HCMD2 dataset, we extracted all the proteins for which a complex structure was available in the PDB, and where at least one experimental partner is also present in the dataset, i.e. proteins corresponding to different chains from the same Protein Data Bank (PDB) ^37^ structure. Any further reference to these proteins uses the PDB code of the experimental structure from which they were extracted, followed by their corresponding one letter chain denomination. For example, chain F of the 1LI1 PDB structure presented in the Results section will be denoted 1LI1_F. Within the 399 proteins originally extracted from the HCMD dataset, only 358 proteins, coming from 138 unique PDB structures, were used for this study after a quality control step, and constitute the *SubHCMD* dataset (listed in Table S1). It is to note that all the proteins are in the bound form. The SubHCMD dataset protein sizes range from 21 to 789 residues, with a median value around 150 residues per protein. In the quality control step, alpha-carbon only structures, proteins with no interaction with other monomers of the same PDB present in the HCMD dataset and proteins with missing docking results were excluded. The large majority of proteins was associated with only one experimental partner but for some proteins over 5 experimental partners were available in the dataset (see Table S1 and Figure S1). Among the 358 proteins of the SubHCMD dataset, 16 proteins are antibody monomers, 6 are antigen monomers, 82 are enzyme monomers, 8 are inhibitor monomers and 246 are classified as « other ».

#### Reduced protein representation

We use a coarse-grain protein model developed by Zacharias ^38^, where each amino acid is represented by one pseudoatom located at the Cα position and either one or two pseudoatoms representing the side-chain (with the exception of Gly). Ala, Ser, Thr, Val, Leu, Ile, Asn, Asp, and Cys have a single pseudoatom located at the geometrical center of the side-chain heavy atoms. For the remaining amino acids, a first pseudoatom is located midway between the Cβ and Cγ atoms, while the second is placed at the geometrical center of the remaining side-chain heavy atoms. This representation, which allows different amino acids to be distinguished from one another, has already proved useful both in protein-protein docking ^38–40^ and protein mechanic studies ^41–43^. Intermolecular interactions between the pseudo-atoms of the Zacharias representation are treated using a soft LJ-type potential with appropriately adjusted parameters for each type of side-chain, see Table I in ^38^. In the case of charged side-chains, electrostatic interactions between net point charges located on the second side-chain pseudoatom were calculated by using a distance-dependent dielectric constant ε=15r, leading to the following equation for the interaction energy of the pseudoatom pair i,j at distance r_ij_: where B_ij_ and C_ij_ are the repulsive and attractive LJ-type parameters respectively, and q_i_ and q_j_ are the charges of the pseudoatoms i and j.

#### Systematic docking simulations

Our systematic rigid body docking algorithm MAXDo (Molecular Association via Cross Docking) was derived from the ATTRACT protocol ^38^ and uses a multiple energy minimization scheme. For each pair of proteins within the SubHCMD dataset, the first molecule (called the receptor) is fixed in space, while the second (termed the ligand) is used as a probe and placed at multiple positions on the surface of the receptor. The initial distance of the probe from the receptor is chosen so that no pair of probe-receptor pseudoatoms comes closer than 6 Å. Starting probe positions are randomly created around the receptor surface with a density of one position per 70 Å^2^. The same protocol is then repeated for the ligand protein. For each pair of receptor/ligand starting positions, different starting orientations were generated by applying 5 rotations of the gamma Euler angle defined with the axis connecting the centers of mass of the 2 proteins (Figure S2a). An extension of the binding site predictions resulting from evolutionary sequence analysis realized with JET ^44^ were used to define the area for docking and restrain the conformational space of the docking algorithm. Thus, only surface regions containing residues predicted to belong to potential binding sites by JET (i.e. residues whose trace value is equal or above 7) will be treated by MAXDo (Figure S2b). For a starting configuration to be treated by the energy minimization scheme, both the receptor and the ligand must present at least one JET predicted residue on their surface that is less than 2 Å away from the axis connecting their centers of mass.

During each energy minimization, the ligand was free to move over the surface of the receptor. Thus, for each couple of proteins P_1_P_2_, considering P_1_ as the receptor and P_2_ as the ligand will lead to substantially the same results than treating P_2_ as the receptor and P_1_ as the ligand (unlike our earlier CC-D studies where the receptor and the ligand proteins were treated differently ^21, 23, 24^). This enables to reduce the number of minimization steps by almost a factor 2.

#### Computational implementation

Each energy minimization for a starting configuration of a pair of proteins typically takes 5 s on a single 2 GHz processor. Between 313 and 2552230 minimizations are needed to probe all possible interaction conformations (with a mean value of 61564 minimizations). Therefore, a CC-D search on the SubHCMD dataset, involving namely = 64261 receptor/ligand pairs, would require several thousand years of computation on a single processor. However, since each minimization is independent of the others, this problem belongs to the « embarrassingly parallel » category and is well adapted to multiprocessor machines, and particularly to grid-computing systems. In the present case, our calculations have been carried out using the *World Community Grid* (WCG, www.worldcommunitygrid.org) during the second phase of the Help Cure Muscular Dystrophy (HCMD2, https://www.worldcommunitygrid.org/research/hcmd2/overview.do?language=en_US) project. It took approximately three years to perform CC-D calculations on the complete HCMD dataset of 2246 proteins. More technical details regarding the execution of the program on WCG can be found in Ref. ^45^.

### Data analysis

#### Definition of surface and interface residues

The relative solvent accessible surface area, calculated with the NACCESS program ^46^, using a 1.4 Å probe, is used to define surface and interface residues. Surface residues have a relative solvent accessible surface area larger than 5 %, whereas interface residues present at least a 10 % decrease of their accessible surface area in the protein bound structure compared to the unbound form.

#### Interface propensities of the surface residues

In order to see whether cross-docking simulations can give us information regarding protein interaction sites, we used the protein interface propensity (PIP) approach ^24^ initially developed by Fernandez-Recio et al. ^15^. The PIP value, representing the probability for residue i of protein P_1_ to belong to an interaction site, is computed by counting the number of docking hits for residue i in protein P_1_, that is, the number of times residue i belongs to a docked interface between P_1_ and all its interaction partners in the benchmark. In earlier works ^21^, we used a Boltzmann weighting factor which would favor docked interfaces with low energies. As a consequence, for a given protein pair P_1_P_2_, all interfaces with a 2.7 kcal.mol^-1^ or more energy difference from the lowest energy docked interface would have a Boltzmann weight lower than 1 % (see ref ^23^ for more details). Here, in order to limit the number of docked interfaces that would have to be reconstructed for determining the interface residues, which is the most time-consuming part of the analysis process, we chose to calculate the residues PIP values using only the lowest energy docking poses within this 2.7 kcal.mol^-1^ criterion, therefore we have where N_pos,P1P2_ is the number of retained docking poses of P_1_ and P_2_ (which will vary with protein P_2_) satisfying the energy criterion, and N_int,P1P2_(i) is the number of these conformations where residue i belongs to the binding interface. Finally, the PIP value for a given residue i belonging to protein P_1_ taking into account the CC-D calculations within the whole benchmark will simply be the average PIP of this residue over all the possible partner proteins P_2_, that is PIP values are comprised between 0 (the residue does not appear in any docked interface) and 1 (the residue is present in every single docked interface involving protein P_1_) and will be used for the prediction of binding sites. For each protein pair in the benchmark, between 1 and 10134 docking poses were kept using the 2.7 kcal.mol^-1^ energy criterion, with an average of 73 docking poses (Figure S3a), and a median value of 26 docking pose kept. These low statistics on each individual protein pair are compensated by the fact that every protein was docked with 358 different partners. Eventually, for each protein in the dataset, between 1293 and 241367 docking poses were used to calculate the residues PIP values, with an average of 25968 docking poses and a median value of 17136 docking poses (Figure S3b).

#### Evaluation of the binding site predictions

Considering the PIP values results for all the residues, we define as predicted interface residues, residues whose PIP value lies above a chosen cutoff. Surface residues can then be divided into 4 classes: true positives (TP) are all the surface residues that are correctly predicted as interface residues, true negatives (TN) being all the surface residues that do not belong to an experimental protein interface and that are predicted as such. False positives (FP) are all surface residues predicted to be in the interface and which do not belong to an experimental interface, and false negatives (FN) being all surface residues that belong to an experimental protein interface but are not predicted as interface residues. We can use the classical notions of sensitivity, specificity and the error function to evaluate their efficiency for the identification of protein interaction sites. Sensitivity (Sen.) is defined as the number of surface residues that are correctly predicted as interface residues (true positives, TP) divided by the total number of experimentally identified interface residues in the set (T). Specificity (Spe.) is defined as the fraction of surface residues that do not belong to an experimental protein interface and that are predicted as such (true negatives, TN). Additional useful notions that are commonly used include the positive predicted value (PPV, also called precision, Prec.), which is the fraction of predicted interface residues that are indeed experimental interface residues (TP/P), the negative predicted value (NPV), which is the fraction of residues that are not predicted to be in the interface and which do not belong to an experimental interface (TN/N) and the false discovery rate (FDR) which is the fraction of residues that are predicted to be in the interface and which do not belong to an experimental interface (FP/P), and corresponds to 1-Prec.

An optimal prediction tool would have all notions (Sen., Spe., Prec. and NPV) equal to unity. If this cannot be achieved, a compromise can be obtained by minimizing a normalized error function based on the sensitivity and specificity values, which is comprised between 0 and 1 and defined as:

Receiver operating characteristics (ROC) curves can be drawn by plotting Sensitivity (also called True Positive Rate) as a function of 1-Specificity (also called False Positive Rate) when changing the PIP value used as a cutoff for prediction. On a classical ROC curve the minimum error corresponds to the point on the curve that is the farthest away from the diagonal (which corresponds to random prediction). The Area Under the specificity-sensitivity ROC Curve (AUC) can be used as a metric to evaluate and compare binding sites prediction performances. Individual ROC curves and individual AUC values can be derived from the study of a single protein, with the TP, TN, FP and FN values being calculated considering only the surface residues of the query protein. For example, an individual ROC curve can be plotted for the 1LI1_F protein, the sensitivity and 1-specifity values being computed on the 177 surface residues of the 1LI1_F protein and the corresponding individual AUC value is equal to 0.360 (BI score without using alternate interfaces). In addition, a general ROC curve and a general AUC value can be computed for the complete SubHCMD dataset, with TP, TN, FP and FN values being processed using all the 50718 surface residues from all proteins in the SubHCMD dataset.

#### Specific case of protein with multiple experimental partners

In a simple case of binding site prediction evaluation, the studied protein presents only one experimental partner, and thus one reference interaction site. The evaluation of the ability of the method to detect this interface then comes down to comparing the predicted interface to the reference experimental one. However, the dataset used in this study includes protein presenting more than one experimental partner and thus more than one interaction site. To evaluate our binding sites predictions while accounting for multiple experimental binding sites, we compared the efficiency of 2 scoring schemes. The first score, called *Global Interface* (GI) score, was obtained by comparing the predicted interface with one single global reference experimental interface generated by concatenating all the available experimental interfaces (Figure S4a). In other terms, if a surface residue is identified as being part of at least one of the experimental binding sites of the target protein, this residue is tagged as interface residue in the global reference experimental interface. Conversely, if a residue is not identified as being part of any of the experimental interfaces of the query protein, it is considered as *non-interface* residue in the global experimental reference. In the second score, named *Best Interface* (BI) score, and similar to the approach developed for the JET^2^ program^35^, the predicted interface is compared to each reference experimental interface separately, and only the predicted interface associated with the best binding site prediction performance was kept (Figure S4b).

#### Interface analysis

##### Search for alternate interfaces protocol

After coloring the residues of the 358 proteins of the SubHCMD dataset according to their PIP value, we realized a systematic visual inspection of the binding site predictions, which led to the identification of alternate binding sites predictions in 188 cases (53% of the protein dataset). To explain the existence of these predicted alternate binding sites, we searched for alternate experimental partners for the 358 proteins of the SubHCMD dataset. In this perspective, we first analyzed for each one of the proteins whether the original PDB structure, from which the protein was extracted, comprises other chains that are not included in the SubHCMD dataset, and which could bind on the predicted alternate interaction site. We distinguish the cases where the alternate experimental partner identified with this analysis is a protein (“Interface with another protein chain of the same pdb structure not included in the dataset”) and the cases where the alternate experimental partner identified consists of a DNA or RNA molecule (“Interface with nucleic acid”). Then, for proteins with predicted alternate binding sites that remained unexplained after this first step of investigation, we searched in the PiQSi (Protein Quaternary Structure investigation) database ^47^ for possible partners fitting our predicted binding sites. This database allows the investigation and curation of quaternary structures by using information about the quaternary structure of homologous proteins. The PiQSi webser thus allowed searching for possible homodimeric structures of the proteins that are not described in the original PDB structure (“Interface from homodimers”) and whose interfaces correspond to the predicted alternate binding sites. We also investigated the quaternary structures of homologous proteins to identify complexes between proteins of the SubHCMD dataset and partners different from those observed in the original PDB structures to explain the predicted alternate binding sites. Again, we differentiate the interface according to the nature of the heterodimer partner identified with PiQSi: “Interface from heterodimers” for proteins and “Interface with nucleic acids” for DNA and RNA.

##### Nucleic acid interfaces

Since 45 proteins in the SubHCMD dataset also present an interface with nucleic acids, we compared the 48 nucleic acid experimental interfaces (NAI) existing between a protein of our dataset and one molecule of nucleic acid (8 with DNA and 40 with RNA, 3 proteins presenting 2 distinct interfaces with RNA) to the 501 initial protein-protein reference experimental interfaces (PPI) formed by proteins of our dataset using the following metrics:

- Fraction of each residue type in the interface (FRI_res(type)_):

where N_res(type)[interf]_ is the number of residues from the corresponding type present in the experimental interface and N_res(total)[interf]_ the total number of residues in the interface.

- Number of each residue type in the interface

- Accessible surface area (ASA) computed with NACCESS ^46^

- Binding site global charge in the interface.

Using a one-tailed Wilcoxon test ^48^, we evaluated for each one of these metrics whether the mean value associated with the NAI was significantly different from the mean value observed in PPI. We also evaluated whether the mean value of the fraction of each residue type in the interface (FRI_res(type)_) in the NAI was significantly different from the one observed in the whole corresponding protein surface (FRI_res(type)[surface]_):

where N_res(type)[surface]_ is the number of residues of the corresponding type present in the total surface and N_res(total)[surface]_ the total number of residues in the surface.

##### Ocupancy rate

For every protein included in our dataset, we computed the occupancy rate of each predicted binding site on its surface, to see if some binding sites are more targeted by the protein partners during the simulations. For a protein pair P_1_P_2_, the occupancy rate of a given binding site on the surface of P_1_ is defined as the fraction of docking poses for which this binding site is selected as the best interface using the BI scoring scheme described earlier. The global occupancy rate for each interface at the surface of P_1_ is then computed as the average of the occupancy rates for all the possible cross-docking partners P_2_.

##### 2P2I inspector

We used the occupancy rates to extract from the SubHCMD dataset 85 proteins (Table S2) presenting one *primary interface* (PrimI), and *one secondary interface* (SecI). To be included in this part of the analysis, proteins should present at least 2 binding sites on their surface, and the occupancy rate for the PrimI should be at least 50% larger than the occupancy rate for the SecI (Table S2). 43 descriptors (Table S3) were computed for each one of the 170 protein complexes (85 complexes forming the PrimI and 85 complexes forming the SecI) using the 2P2I inspector website ^49^. Predicted interfaces with no corresponding experimental binding site or interfaces with a non-protein partner (nucleic acids) could not be included in this part of the study since the 2P2I inspector website only accepts protein PDB files as input. We then evaluated for each descriptor if the mean value observed for this descriptor in the PrimI was significantly different from the one observed for the SecI using a one-tailed paired Student test.

#### Statistical analysis

The R Wilcoxon Mann-Whitney algorithm ^50^ was used to compute the one-tailed Wilcoxon test to evaluate whether the mean value observed for the fraction of each residue type in the interface, the number of each residue type in the interface, the accessible surface area and the binding site global charge in the NAI were significantly inferior (option “less”) or significantly superior (option “greater”) than the mean values observed in the PPI. To correct for multiple tests for the fraction of each residue type in the interface and the number of each residue type in the interface, the Bonferroni threshold ^51^ was applied to evaluate the statistical significance of the p-values obtained (Bonferroni threshold is equal to 0.05/N_res(type)_, with N_res(type)_ being the number of residue type tested).

The R tool was used to compute the one-tailed paired Student test ^52^ to evaluate for each 2P2I inspector descriptor if the mean value observed for this descriptor in the PrimI was significantly inferior (option “less”) or significantly superior (option “greater”) than the one observed for the SecI. To correct for multiple tests, the Bonferroni ^51^ threshold was applied to evaluate the statistical significance of the p-value (Bonferroni threshold is equal to 0.05/ N_descriptors_, with N_descriptors_ being the number of computed 2P2I inspector descriptors).

All graphics were produced using the statistical and graphical tool R (http://www.r-project.org/).

## Results

We must recall that, since the point of this work is to investigate the prediction of binding sites on protein surfaces with no prior knowledge of the binding partners, and not the correct docking of experimentally known partners, which can be achieved via other more effective but much more computationally demanding methods ^53^, we did not evaluate the quality of the best structural predictions for the docked complexes. However, in an earlier work ^21^, where we performed cross-docking simulations on a limited test-set involving 12 proteins (using their bound structures), our method was able to predict correctly the position of the ligand protein with respect to its receptor with an rsmd of the Cα pseudoatoms below 3 Å, thus validating the quality of the force-field used in our systematic rigid body docking algorithm called MAXDo. Furthermore, this force-field, which was originally developed by Zacharias for protein-protein docking ^38^, has been successfully used on numerous occasions for the prediction of protein complex structures, especially during the CAPRI contest where the unbound structures of the protein partners are used ^39, 54, 55^.

### Identification of protein interaction sites

Classically, the performance of a cross docking method to identify protein interaction sites is assessed using complexes formed by experimental partners. Therefore, the evaluation of the method is limited to the prediction efficiency for identifying one reference experimental interaction site on each protein’s surface. However, many proteins are known to be able to interact with different partners and to present multiple binding sites on their surface ^56–58^. Among the 358 proteins used for this CC-D study, 96 present more than one known experimental partner and thus potentially complex binding sites distributions (with single sites binding several partners or/and multiple patches, see ref. ^36^). Consequently, we developed a new protocol for the evaluation of the binding site predictions for these proteins and we compared the use of two scoring schemes accounting for multiple binding sites, the *Global Interface* (GI) score and the *Best Interface* score (BI) (detailed in the Material and Methods section), for evaluating the binding sites prediction.

The performances of the GI and BI scores are summarized in Table 1, Figures 1 and 2. Differences only appear for the 96 proteins (out of the 358 proteins included in the dataset) associated with more than one experimental partner, the two scoring schemes being identical when only one experimental partner is available (Figure S5). When considering the complete dataset of 358 proteins, the BI score is associated with a higher overall AUC value (0.732 compared to 0.696, while random predictions would give an AUC value of 0.5) and lower minimum error (0.48 compared to 0.51) than the GI score (Figure 1). For 80 over the 96 proteins with more than one experimental partner included in the dataset, the best individual AUC is obtained with the BI scoring scheme. The GI scoring scheme is associated with an individual AUC superior to the BI scoring scheme for only 12 proteins over 96. As a consequence, we chose to use the BI score for the rest of the analysis.

**Figure 1.**
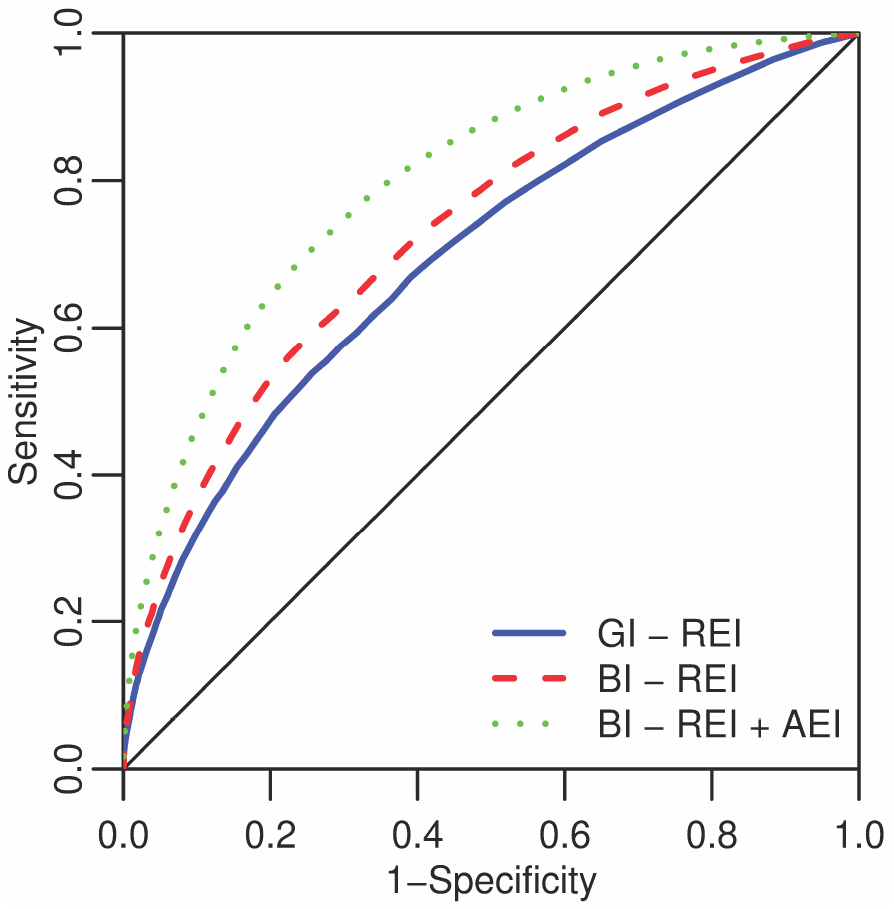
Overall ROC curves of the PIP prediction obtained using the *Global Interface* score using only the reference experimental interfaces (REI) (blue line), using the *Best Interface* score with the REI (red line), and using the BI score with both the REI and the alternate experimental interfaces (AEI) identified with other protein partners or with nucleic acid molecules (green line). The diagonal dotted line corresponds to random prediction. See Figure S4 for a schematic description of how the BI and GI scores are calculated.

**Figure 2.**
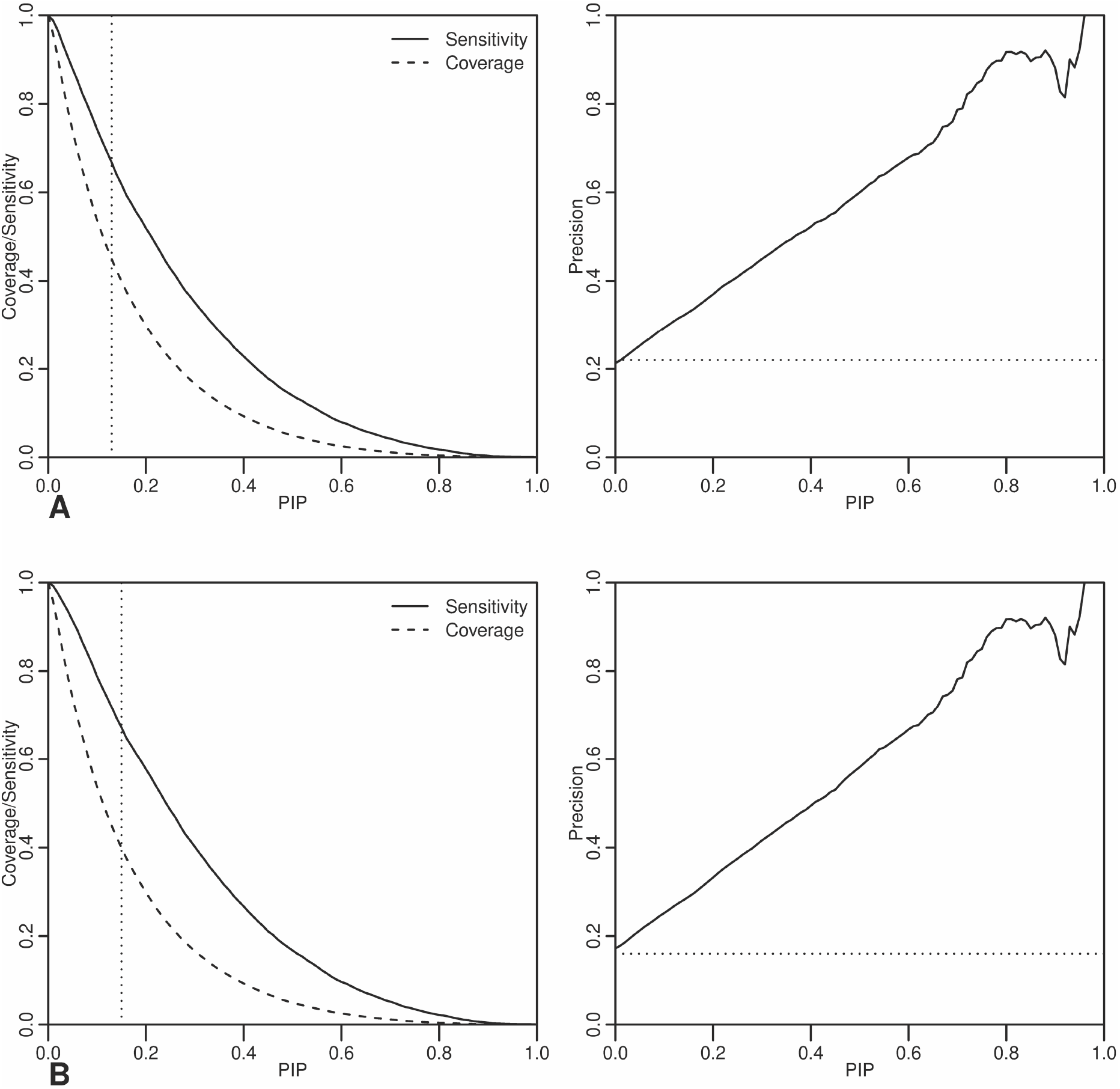
Enrichment of the interface residues from the 358 proteins of our CC-D dataset using the PIP index shown by comparing the fraction of true interface residues detected (sensitivity, solid line) with the total fraction of residues detected (coverage, dashed line) as a function of the PIP cutoff using the GI score (a) or the BI score (b). The vertical dotted lines correspond to the position of the optimal PIP cutoff leading to the minimal error function. Precision as a function of the PIP cutoff, the dotted horizontal line corresponds to random predictions: (c) GI score, (d) BI score)

### Identification of alternate interfaces different from the reference experimental interfaces

#### Predicted alternate interfaces

The efficiency of the PIP values for predicting protein interfaces in different protein functional groups has been assessed in our earlier work ^24^ on 168 proteins from the Docking Benchmark 2.0 ^28^. Here the individual AUC values obtained with the 358 proteins of the dataset show a large distribution in the binding site prediction efficiency depending on the protein studied, even when using the BI score (see Figure S6). 66 proteins are associated with very high binding site prediction efficiency, with individual AUC values superior to 0.9, whereas the individual AUC values lie below the random prediction threshold of 0.5 for 68 proteins (i.e. 19% of the complete set). By coloring the protein surface residues with the PIP values resulting from the cross-docking calculations, we could observe predicted alternate interfaces that are visibly different from the expected reference experimental interfaces for 188 out of the 358 proteins included in the dataset (53%). This visual observation can be partially correlated with the individual AUC values since for all 68 proteins for which the binding site predictions performance are below the random prediction threshold (AUC equal to 0.5), predicted alternate interfaces are observed. Conversely, only 24 % of proteins with individual AUC values above 0.75 present predicted alternate interfaces, and this rate decreases to 9 % when focusing on proteins with individual AUC values above 0.9. In order to proceed in a more systematic and observer-independent approach, we used the correlation between the visual observations and the false discovery rate (FDR) computed using the optimal PIP-cutoff of 0.15 (Figure S7). We computed the optimal threshold of FDR with the normalized error function, and obtained an optimal value of 0.67. Using this value, we assume that a protein associated with a FDR superior to 0.67 will present an alternate binding site whereas a protein with a FDR inferior to 0.67 will not present a predicted alternate binding site. Indeed, 164 proteins over the 188 for which predicted alternate interface were visualized (87%) present a FDR superior to 0.67 and only 16% of proteins for which no predicted alternate interface could be observed present a FDR superior to the optimal FDR threshold. Binding site predictions for proteins with no visual predicted alternate interface and FDR values superior to 0.67 include the reference experimental interfaces, but are larger than them.

#### Rationalizing the prediction of alternate interfaces

In an earlier study ^24^, we showed for some examples that the apparent failure of cross-docking calculations for predicting experimental binding sites could be explained by the detection of alternate interfaces formed by the protein with other biomolecular partners that are not present in the original reference dataset. We decided to push this analysis further in an exhaustive fashion by investigating whether it is possible to find experimental partners that would bind on the predicted alternate interfaces and to which extent. For 146 proteins over the 188 proteins (78%) with predicted alternate interfaces, we could identify an alternate experimental partner, not included in the original dataset (and therefore in the docking calculations), whose experimental interface with the protein under study corresponds to the predicted alternate interface. Eventually, predicted alternate interfaces could be segregated in 3 categories: interfaces with another protein partner (interfaces with another protein chain from the same PDB structure not included in the dataset, interfaces from homodimers; interfaces from heterodimers); interfaces with nucleic acids; unexplained predicted alternate interfaces (note that small ligand-binding sites, as described in ref. ^35^ were not detected in the process).

##### Interfaces with another protein partner

*Interface with another protein chain from the same PDB structure not included in the dataset.* Figure 3a presents the case of the structural protein collagen alpha 2 (IV) (1LI1_F). In our dataset, 2 experimental partners for this protein are included: the collagen alpha 1(IV) (1LI1_B) and a second monomer of collagen alpha 2 (IV) (1LI1_C) (shown in grey and black, respectively, on Figure 3a). However, the residues of 1LI1_F presenting the highest PIP values do not belong to any of the expected reference experimental interfaces, leading to a low individual AUC value of 0.360. Nevertheless, the global 1LI1 PDB structure comprises six protein chains in total, and the residues of 1LI1_F presenting the highest PIP values are in fact those involved in the interface with two other chains of the same PDB structure not included in our dataset, 1LI1_D (Figure 3a, shown in smudge green) and 1LI1_E (Figure 3a, shown in green). When adding the information about these new reference experimental interfaces, the individual AUC of the binding site prediction associated with 1LI1_F increases to 0.689 (corresponding to the interface with 1LI1_E). One must note that the binding sites of 1LI1_D and 1LI1_E on the surface of 1LI1_F are contiguous. These two sites form a large patch that can bind multiple partners (similar to what could be observed in refs. ^35^ and ^36^), and in this specific case, the GI score computed by comparing the PIP values to the global reference interface obtained by concatenating the 1LI1_D and 1LI1_E experimental interfaces performs better than the BI score (0.800 vs 0.689). We proceeded in the same way for each one of the proteins of our dataset and we identified 81 proteins over 188 with predicted alternate interfaces (43%), whose predicted alternate binding site corresponds to the interface with another protein chain of the same PDB that was not included in the original dataset. In addition to these 81 proteins, 10 proteins (1AVO_B, 1D8D_A, 1D8D_B, 1JJO_C, 2NNA_A, 2NNA_B, 3BRT_B, 3BRT_C, 3C5J_A, 3C5J_B) present enhanced binding site predictions when adding new reference experimental interfaces from other protein chains of the same PDB structure not included in the dataset. These 10 proteins do not present obvious predicted alternate interface, i.e. their individual AUC values are superior to 0.5, their FDR values computed at the optimal PIP cut-off of 0.15 are below 0.67 and no predicted alternate interface could be detected by visual inspection (Figure S8a-j, left panels). In fact, as illustrated in Figure S8a-j (right panels), the new experimental interfaces are overlapping the initial reference experimental interface, explaining why no predicted alternate interface was detected at first. Finally, taking into account all these new reference experimental interfaces, the global performance of the binding site prediction obtained with the BI score and measured using the overall AUC increases from 0.732 (value obtained with the initial reference experimental interfaces) to 0.776 (Figure S9, green line).

**Figure 3.**
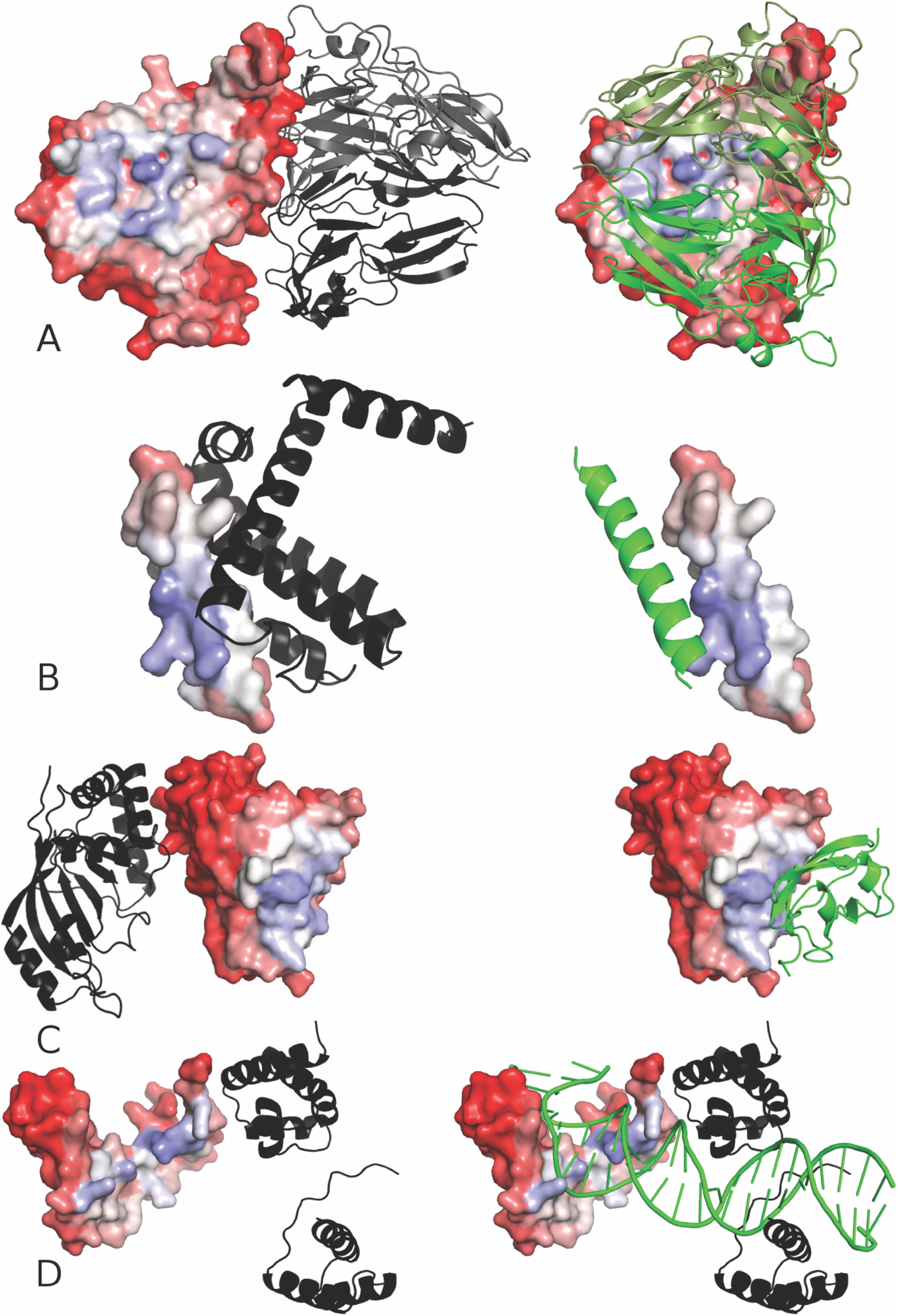
Mapping the PIP values on a protein’s surface, high PIP residues are shown in blue and low PIP residues are shown in red. The reference experimental partner (ref) is shown in a black or grey cartoon representation, while the alternate partner (alt) is shown in green. (a) The collagen alpha 2 (IV) (1LI1_F) with collagen alpha 1 (IV) (1LI1_B (ref) in grey), and other collagen alpha 2 (IV) monomers (1LI1_C (ref) in black, 1LI1_D (alt) in smudge green and 1LI1_E (alt) in green). (b) Beclin 1 (2P1L_B) with Bcl-X (2P1L_A(ref)) and another monomer of Beclin 1 identified using the PiQSi webser (alt). (c) RalA protein (2BOV_A) with C3 ribosyltransferase (2BOV_B(ref)) and Sec5 protein (1UAD_C(alt)). (d) Transcription factor SOX-2 (1GT0_D) with the octamer-binding transcription factor 1 (1GT0_C(ref)) and DNA (1GT0_A/B(alt)).

*Interfaces from homodimers and heterodimers.* Using the PiQSi webserver ^47^, we searched for alternate partners for the 188 proteins with predicted alternate binding sites. For 12 proteins over these 188 proteins (6%), interfaces formed in homodimeric structures (described in the PiQSi database) overlap alternate binding sites predicted using the PIP values, as illustrated in the Figure 3b. Adding these new reference experimental interfaces to the binding site prediction calculations leads to only a small increase of overall AUC (0.736 vs 0.732, Figure S9, blue line). In the same way, for 22 proteins over the 188 proteins investigated (12%), we could identify homolog proteins forming heterodimeric complexes that could explain the predicted alternate binding site. This is notably the case for the RalA protein (2BOV_A) that presents one of the lowest binding site predictions quality (individual AUC = 0.232). As shown in the Figure 3c, cross-docking led to the prediction of an alternate binding site on the opposite side of 2BOV_A compared to the reference experimental interface with the C3 ribosyltransferase (2BOV_B). Using the PiQSi database, we could identify another complex (PDB ID: 1UAD) between the same RalA protein (1UAD_A) and a different partner, the Sec5 protein (1UAD_C). The experimental interface between RalA and Sec5 perfectly concurs with the alternate interface predicted by cross-docking on the surface of RalA (2BOV_A). The inclusion of these heterodimeric experimental interfaces in the reference binding sites dataset leads to an increase of binding site prediction performance using the BI score, with an overall AUC value of 0.749 (Figure S9, cyan line).

#### Single docking vs Cross docking

To study whether similar interface prediction could be obtained by docking a single, randomly selected in the PDB, protein against one protein of the subHCMD dataset, i.e. using *single-docking*, we used the 4 proteins previously presented as test cases: 1LI1_F, 2P1L_B, 2BOV_A and 1GT0_D. For each case, we computed all the individual AUC values obtained when performing single-docking with every proteic partner from the SubHCMD dataset (Figure S10).

#### Correlation between JET score and PIP values

To reduce the computational time associated with cross-docking, we used binding sites predictions from JET to restrain the conformational space of the docking algorithm. The JET score we used, ranging from 0 to 10, represents the number of time a residue was selected as interface residue by an iteration of iJET. To explore whether this initial pre-filtering protocol impacts the resulting PIP values and thus the binding site predictions, we evaluated the correlation between <PIP> (i.e., the average PIP value of a given residue after docking with all the proteic partners from de SubHCMD dataset) and JET score values for each residue of each protein (see Figure S11 for examples of correlation between <PIP> and JET score within a protein). An analysis of variance (ANOVA) on the distribution of <PIP> as a function of the JET score (Figure 4) shows that the average <PIP> for each JET score group differ significantly (p-value below 2E^-16^). This is particularly true for the two extreme values of JET score, 0 and 10, associated with respectively lower and higher average <PIP> compared to the other JET score groups. However, an ANOVA on the distribution of <PIP> for JET score groups equal to 5, 6, 7, 8 and 9 results in a p-value of 0.126. The average <PIP> is not significantly different between these groups, even though only residues with JET score equal or above 7 were considered as predicted interface residues in our protocol. We assume that, beside the agreement for binding site predictions between JET score and <PIP> values, the cross-docking approach is not biased by the pre-filtering step using JET score. Residues not predicted as interface residues using JET could be identified using the cross-docking approach and conversely, residues predicted as interface residues by JET were associated with low <PIP> values. One such example is the case of the protein 3BES_R (Figure S12), for which accurate binding site predictions could be obtained both by using JET scores and cross-docking results, except that both approaches are pointing at two different interfaces corresponding to different partners.

**Figure 4.**
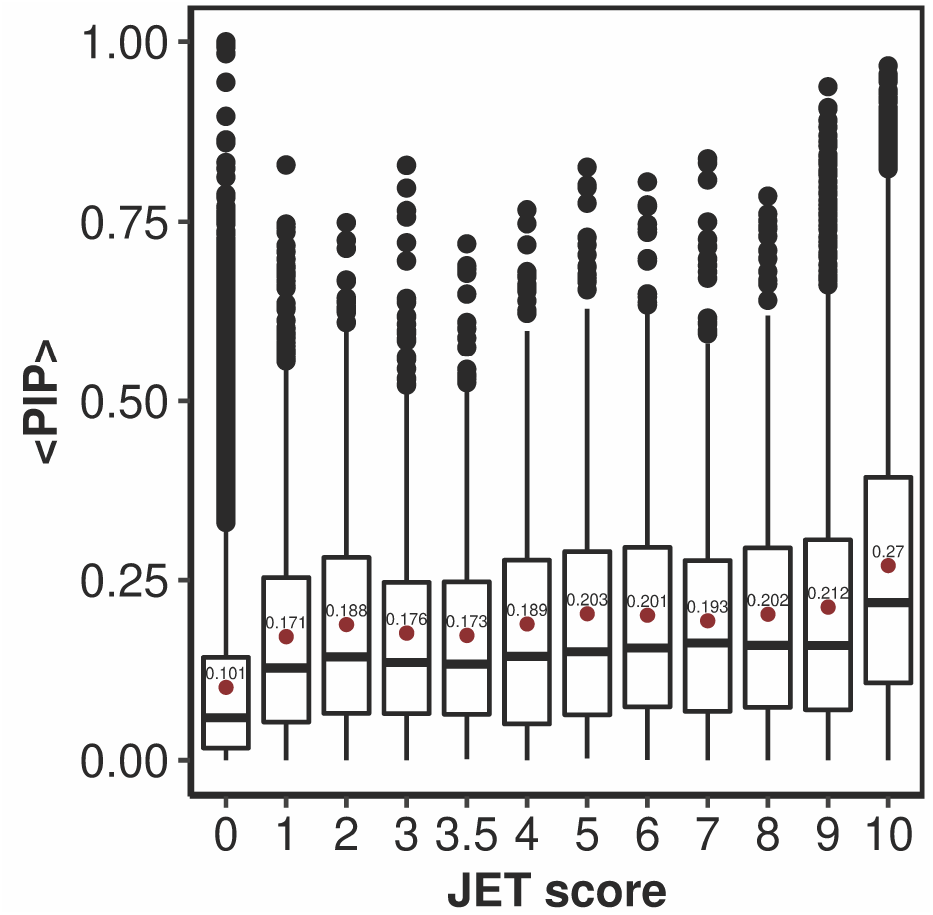
Boxplot of the distribution of <PIP> values as a function of JET score for the complete SubHCMD dataset. The red dot and its corresponding label indicate the mean value of <PIP> for each JET score.

##### Interfaces with nucleic acid molecules

For 31 proteins over the 188 with predicted alternate interfaces (16%), the binding signal observed with cross-docking simulations turned out to correspond to a binding site with nucleic acids. The transcription factor SOX-2 (1GT0_D) is an example of such proteins. The PDB structure 1GT0 represents the complex formed between SOX-2 (1GT0_D), the octamer-binding transcription factor 1 (1GT0_C) and a DNA double strand. The 1GT0_C and 1GT0_D proteins were included in our cross-docking dataset, unlike the DNA double strand. When focusing on the cross-docking binding site predictions for 1GT0_D (Figure 3d), a weak binding site prediction of the interface with 1GT0_C is obtained (individual AUC of 0.595). Indeed, the residues of 1GT0_D presenting the highest PIP values are those surrounding the DNA double strand. Thus, using our cross-docking procedure, we could identify interfaces with non-proteic alternate partners, like nucleic acids, not included in the original dataset used to run the simulations. New binding site prediction performances were computed including this additional experimental interfaces with nucleic acids leading to a slight increase in the overall AUC value (0.739 vs 0.732) (Figure S9, magenta line).

Since our docking scheme has been developed to study protein-protein interactions and only proteins were included in the CC-D simulations, its ability to predict nucleic acid binding sites came as a surprise to us. However, these predictions concur with earlier work using evolutionary data, which showed that DNA and RNA binding sites do overlap with protein binding sites in many cases ^44^. We thus decided to analyze the nucleic acid interfaces (NAI) at our disposal to try to understand this point (Figure S13). We compared the mean value of the fraction of each residue type in the interface (FRIres(type)), of the number of each residue type in the interface, of the accessible surface area and of the binding site global charge in the interface in the NAI and PPI using a one-tailed Wilcoxon test. We also compared the FRIres(type) in the interface in the NAI compared to the fraction of each residue type observed in the whole corresponding protein surface (FRIres(type)[surface]) using a one-tailed Wilcoxon test. Compared to PPI, our data show a global enrichment of the NAI in the following residues: Arg, Lys (both residues are positively charged in the Zacharias coarse-grain model), His, Gly, and Ser; while NAI are poorer in Asp and Glu (negatively charged in the Zacharias coarse-grain model), Phe and Val. In addition, NAIs appear to be significantly more charged than PPIs (mean binding site global charge of 9.5, vs 0.075 with p-values inferior to 2.2E^-16^) but no significant difference in size is observed (with a mean ASA of 2324 Å^2^ vs 1956 Å^2^). We also evaluated whether the nucleic acid binding predictions obtained with the cross-docking results were biased by the JET binding site predictions by again using an ANOVA on the <PIP> distribution for all the residues predicted to belong to nucleic acid binding site as a function of their JET score (Figure S14). The mean <PIP> value is significantly different between the JET score groups (p-value < 2E^-16^), since the mean <PIP> value is lower in the group of JET score equal to 0 and higher in the group of JET score equal to 10. However, if we reiterate the analysis by excluding these two JET score groups, we obtain a non-significant p-value equal to 0.601 showing that the mean <PIP> value is not significantly different between JET score groups corresponding to residues not predicted as interface residues by JET (JET score equal to 1, 2, 3, 4, 5 and 6) and JET score groups corresponding to residues predicted as interface residues by JET (JET score equal to 7, 8 and 9).

##### Unexplained interfaces and global prediction performances

For only 42 proteins over the 188 (22%) with a predicted alternate interface, we could not identify an alternate biomolecular partner explaining the predicted alternate binding site. When adding all the newly identified alternate reference experimental interfaces to the initial ones, the binding sites prediction performances jumped from 0.732 to 0.801 (Figure 1, green line and Table 1), with enhanced sensitivity, specificity and precision values. Using the BI score with reference experimental PPIs for the original dataset, we failed to correctly predict the binding site location on the protein surface for 68 cases (out of 358 proteins), which presented an individual AUC value below 0.5. After including the alternate experimental binding sites in our calculations, only 10 proteins kept an individual AUC value below 0.5 (see Figure 5).

**Figure 5.**
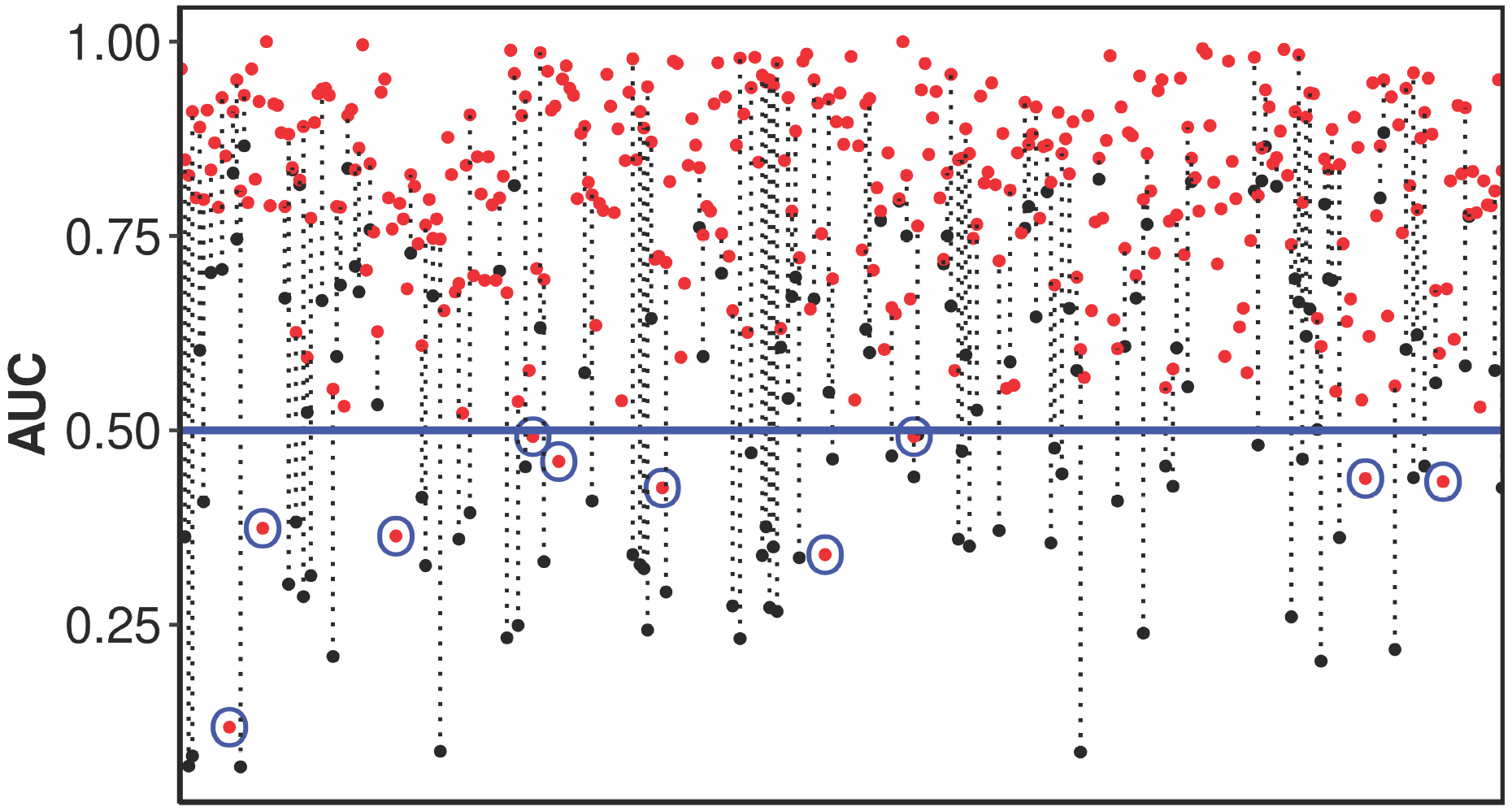
Individual AUC value for each protein in the CC-D dataset obtained with the BI score. The black dots represent the individual AUC values obtained with the reference experimental interfaces (REI). The red dots represent the individual AUC values obtained when including the alternate experimental interfaces (AEI) identified with other protein partners or with nucleic acid molecules to the reference experimental interfaces. The blue circles highlight the 10 proteins that present individual AUC values below the random threshold of 0.5 (blue line) after including the alternate experimental interfaces.

#### Are some interfaces more targeted than others with the cross-docking simulations?

The binding site predictions resulting from cross-docking simulations were used to calculate occupancy rate for each binding site (Figure S15); i.e. the fraction of docking poses for which a given binding site is selected as the best interface (see the Material and Methods section). The analysis of the occupancy rate distribution leads to the distinction of two major behaviors. The first concerns proteins with one binding site that is predominantly targeted by the different partners during the cross-docking simulations (Figure S16a) and the second concerns proteins presenting similar occupancy rates for their different predicted binding sites (Figure S16b). To gain insight into why some binding sites prevail on others, we used the 2P2I inspector v2.0 website ^27^ to compute 43 descriptors for the primary (PrimI) and secondary (SecI) interfaces found on 85 proteins (Table S2 and Figure S17). For each of the 2P2I inspector descriptors, we performed a statistical analysis using a one-tailed Student test to evaluate whether this descriptor’s mean value is significantly inferior or superior in the PrimI compared to its mean value in the SecI. To correct for multiple tests, the Bonferroni threshold was applied to evaluate the statistical significance of the p-value. The percentage of polar contribution to the interface is the only descriptor presenting a mean value significantly inferior in PrimI compared to SecI (Figure S18 and Table S3). The mean values of 15 descriptors within the PrimI are significantly superior to their mean value associated with the SecI (Figure S19 and Table S3). These descriptors belong to the following categories: *Number of non-bounded contacts* (ContRes, ContRN, ContRP, ContRHyd, ContRC, Nb of non-bonded contacts), *Gap volume*, *Accessible surface area* (% Interface Accessible Surface Area, Interface Accessible Surface Area, % Non polar contribution, Total Interface Area), *Segments* (Number of Segments, Total Nb of Segments, a segment being defined as a stretch of interface residues that may contain non-interface residues but not more than four consecutive ones), and *General Properties* (Planarity).

## DISCUSSION

### Identification of protein interaction sites

The ability of cross-docking simulations to identify binding sites on protein surface has already been proved and studied with docking benchmark datasets of 12 proteins for the proof of concept ^21^, and 168 proteins for more recent studies ^23, 24^. However, to the best of our knowledge, the impact of the presence of multiple binding sites on proteins surface, a common feature observed in the protein world, has never been systematically addressed. We thus decided to tackle this problem, using 358 non-redundant proteins that were not extracted from a docking benchmark, but from a database of proteins conceived to study proteins involved in neuromuscular diseases or in the pathways monitoring essential cardiac or cerebral mechanisms. Among those proteins, 96 presented more than one experimental partner in the dataset and thus more than one binding site on their surface. We compared the use of 2 scores, GI and BI to ensure the optimal evaluation of binding sites predictions. Using the GI scoring scheme with proteins presenting multiple binding sites on their surface, the performance of the cross-docking method to make accurate binding site predictions could appear artificially low in cases where one binding site was correctly predicted and the other(s) only partially. The BI score, which only uses as reference the experimental interface the best predicted, avoids this bias and we thus decided to perform the rest of the study with the BI score. Note that this *best patch* approach was also used to compute the performances for predicting protein binding sites with the JET^2^ tool and with similar conclusions ^35^. We obtained good overall binding site prediction performances with the BI score that are only slightly inferior to the ones obtained in earlier cross-docking studies involving respectively 12 ^21^ and 168 proteins ^23^. The differences observed in terms of specificity and sensitivity could be explained by the differences in the proteins dataset used. In the two previously published studies, proteins were extracted from docking benchmark dedicated to protein docking evaluation, whereas in this study, the proteins were selected according to their biological functions and not for benchmarking purpose, and thus represent more challenging cases for docking predictions. In addition, one should note that the SubHCMD dataset only comprises bound protein structures, and our previous studies ^24^ have shown that these will perform less well than unbound structures for binding site prediction via CC-D, probably because a protein’s specific conformational adaptation to a given partner might decrease the quality of its binding to other potential partners. We chose to use the SubHCMD dataset and not a benchmark database since the SubHCMD is part of a bigger dataset we are currently investigating to learn more about some proteins involved in neuromuscular diseases. However, a comparable study could have been achieved with a benchmark database such as the Docking benchmark 5.0 ^25^, DOCKGROUND ^26^, or 2P2IDB ^27^. It is to note that some proteins included in our SubHCMD dataset are also present in these benchmarking databases (1ATN_A, 1ATN_D, 1US7_A, 1US7_B, 2C0L_A, 2C0L_B, 2OT3_A and 2OT3_B are in both SubHCMD and Docking Benchmark 5.0; 1AOX_A, 2BOV_A, 2BOV_B, 1M63_E, 2C0L_A, 2C0L_B, 1MHW_A, 3BS5_A and 3BS5_B are in both SubHCMD and DOCKGROUND).

### Identification of alternate interfaces different from the reference experimental interfaces

Even using the BI score, i.e. the score accounting the best for multiple binding sites, the quality of prediction associated with some proteins was very low. A systematic visual inspection of the proteins, realized by coloring the proteins residues according to their PIP values, highlighted 188 cases where the predictions led to the identification of alternate binding sites. We could define a threshold of FDR (computed at the optimal PIP cut-off of 0.15) equal to 0.67 that could be used to replace the systematic visual inspection to detect proteins with predicted alternate binding site. For 78% of the proteins for which we could visualize a predicted alternate binding site, we identified an alternate partner, not included in the SubHCMD dataset, which fits the corresponding interface. For most cases, this alternate partner is a protein, but more surprisingly, the alternate partner can also consist of nucleic acids. Since our docking protocol was tuned for protein-protein docking and only protein structures were included in the calculations, our ability to predict nucleic acid binding site was unexpected. We could highlight differences between NAI and PPI in terms of amino acid composition, particularly for charged residues, positively charged residues being favoured in NAI and negatively charged residues being favoured in PPI, in agreement with earlier studies (see for examples ^59–63^). The NAI also presented higher global charge and ASA values. The higher global charge observed in NAI (linked to the higher abundance of positively charged residues) could be an explanation of the ability of our CC-D protocol to identify NAI, since we used the first version of the Zacharias coarse-grain model’s force field that overestimates the contribution of the electrostatic term to the overall interaction energy. Even if JET is able to identify DNA and RNA binding sites ^44^, we did not identify an overall correlation between <PIP> and JET score values for residues in proteins presenting nucleic acid binding sites, even if the mean value of <PIP> was the lowest in the JET score=0 group, and the highest in the JET score=10 group. Thus, the identification of nucleic acid binding sites cannot be explained only by the JET binding site predictions pre-filtering. Finally, for only 22% of the 188 proteins presenting an alternate predicted binding site, we could not find an experimental partner binding on the alternate interaction site, but we cannot rule out the fact that the current available experimental data are yet insufficient. For all the proteins for which alternate partners were found, taking into account all the experimental interfaces residues (both from the reference experimental interface and the alternate experimental interface) for the evaluation of the binding site predictions via cross-docking leads to a significant increase of the overall AUC (from 0.732 to 0.801), and of the method’s sensitivity, specificity and precision, that were similar to those obtained in previous studies ^21, 24^. Furthermore, for only 10 proteins the binding site predictions were associated with individual AUC values below the random threshold of 0.5, which represents a significant improvement compared to the case where only reference experimental interfaces are taken into account (68 proteins with individual AUC values < 0.5). For a protein with multiple binding sites on its surface, one binding site (which we defined as “PrimI”) could be targeted preferentially compared to the others (called “SecI”). To understand why some interfaces were more targeted than others during our docking protocol, we used structural descriptors to compared both type of interfaces and we showed that the “PrimI” were larger, more hydrophobic, more planar and characterized by a larger number of contacts between the 2 partners than the “SecI”. Interestingly, these findings give some insights regarding structural determinants preferences in protein-protein interactions linked to our docking protocol, and can be compared with the structural features characterizing protein-protein interaction sites identified in previous studies ^64–68^. According to the protein-protein interfaces size classification established by Lo Conte et al. ^65^, “PrimI” size median value is typical of the *large* interface category (3359.5 Å²) whereas the “SecI” size median value (1349.3 Å²) is similar to the sizes observed in *the standard-size* interfaces. Moreover, De et al. ^68^ showed that nonobligatory protein-protein interfaces, i.e. interfaces observed in transient protein-protein complexes, present generally smaller interface areas associated with weaker interactions than the obligatory protein-protein interfaces (i.e. permanent complexes interfaces). Thus, the ability of our docking protocol to preferentially predict large protein-protein interfaces makes it relevant for the determination of permanent interfaces. Focusing on the hydrophobicity parameter, different trends were observed in the studies, some interfaces being more hydrophobic (homodimers ^67^, standard-size protease-inhibitor interfaces ^65^) and other relatively polar (antibody-antigen interfaces ^65^). ^64^ Finally, the relative flatness of protein-protein interfaces has already been described ^64^, Chakrabati and Janin ^66^ however, found that this flatness was variable. For example antibody-antigen interfaces are more planar than average, whereas protease-inhibitor interfaces are less planar than average. Finally, we showed that the cross-docking protocol is essential for accurate binding site predictions by comparing the results of single docking and cross-docking calculations. If one selects a single random protein as a partner for single docking, one may fail to correctly predict interface residues, and only the large number of proteins used during the cross-docking simulations can ensure the statistical robustness of the predictions. Additionally, a previous cross-docking experiment conducted with a different docking algorithm ^69^ was analyzed to explore possible binding site predictions discrepancies between docking of experimental partners and docking of non-interactors. The results showed that in some cases the individual partner preferentially targets the experimental binding site whereas the non-interactors bind to another location, but in the majority of cases, both the individual partner and the non-interactors target the same site. We also show that using the JET score to pre-filter the starting conformations of the docking did not bias the binding site predictions obtained with the cross-docking calculations. It is to note that in our protocol, the threshold of the JET score we used (corresponding to the number of time a residue was predicted as interface residue by an iteration of iJET) was defined as 7. However, lower values of JET score can still be meaningful for binding site predictions, as mentioned in the ref. ^44^.

## Conclusion

In this work, we evaluate the ability of our cross-docking protocol to lead to the identification of interaction sites on proteins surfaces by using the PIP index computed for each protein residue. Unlike previous studies, the SubHCMD dataset of 358 proteins used is not a benchmarking dataset developed for protein docking evaluation, but a subgroup from a larger database (2246 proteins) built around about 400 proteins known to be involved in neuromuscular diseases. The SubHCMD proteins thus represent more challenging test cases for binding sites prediction, particularly for those presenting multiple binding sites on their surface. We first compared the use of two scores accounting for proteins with multiple binding sites: the GI score that compares PIP values to a single global reference experimental interface and the BI score that treats separately each known experimental interface and keeps only the one that is best predicted. Overall, high quality binding sites predictions are obtained using the BI score, which is best suited for proteins presenting multiple binding sites. However, some proteins were associated with very low performance. In particular, 68 proteins (19 % of the complete set) presented an individual AUC below the random threshold of 0.5. For these 68 proteins and 120 more, alternate interfaces different from the expected reference experimental interfaces were predicted. For about 80% of them, we could identify hidden partners, i.e. a partner not included in the SubHCMD dataset and that binds in the predicted interaction site. An unexpected result was the prediction of nucleic acid binding sites, probably linked to the force field used, which overestimates the electrostatic interactions between charged residues. The ability of the cross-docking simulations to guide towards the identification of these alternate binding sites (and thus of alternate potential partners) is a very promising result since the final aim of the project is to understand the function, and to identify binding sites and potential binding partners for the 400 proteins involved in neuromuscular diseases included in the HCMD (Help Cure Muscular Dystrophy) dataset. We also showed that for proteins with multiple binding sites on their surface, one binding site can attract the majority of the docking partners, and we could identify some interface structural determinants linked to our docking protocol. This study demonstrates that our cross-docking simulations protocol enables to identify accurately and efficiently binding sites on proteins surfaces and defines a first step in the analysis of the whole HCMD dataset of 2246 proteins.

## Acknowledgments

This work was originally carried out in the framework of the DECRYPTHON Project, set up by the CNRS (Centre National de la Recherche Scientifique), the AFM (French Muscular Distrophy Association) and IBM. The cross-docking calculations were carried out on the *World Community Grid* (www.worldcommunitygrid.org) in phase 2 of the Help Cure Muscular Distrophy project. The authors wish to thank all the members of the WCG team for adapting our docking program for use on this grid, and also WCG volunteers for donating the computing time that made this work possible. S. S.-M., A. C. and N. L. thank the Mapping Project (Investissement d’Avenir) ANR-11-BINF-003 for postdoctoral funding. A.C. thanks the Institut Universitaire de France.

This work was supported by the “Initiative d’Excellence” program from the French State (grant « Dynamo », ANR-11-LABX-0011).

## Supporting Information

Supplementary data to this article can be found online.

Binding site predictions on the SubHCMD Protein Benchmark resulting from cross-docking simulations can be downloaded as an archive using the following link: https://owncloud.galaxy.ibpc.fr/owncloud/index.php/s/kjcgtOJ2GggvIhh

