## Supplementary Materials for "Hidden partners: Using cross-docking calculations to predict binding sites for proteins with multiple interactions"

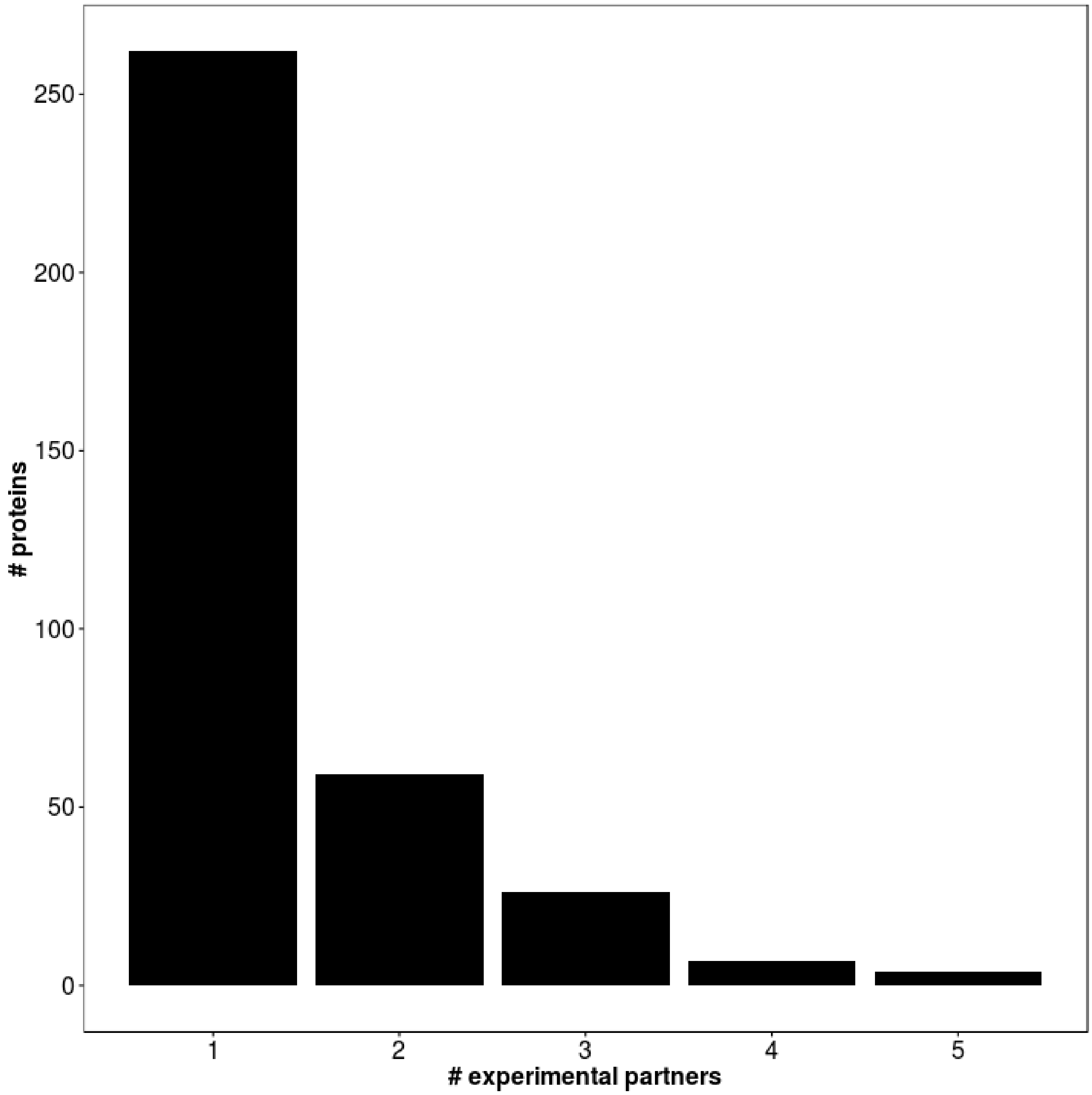


**Figure S1:** Distribution of the number of experimental partners among the proteins included in our cross-docking dataset.

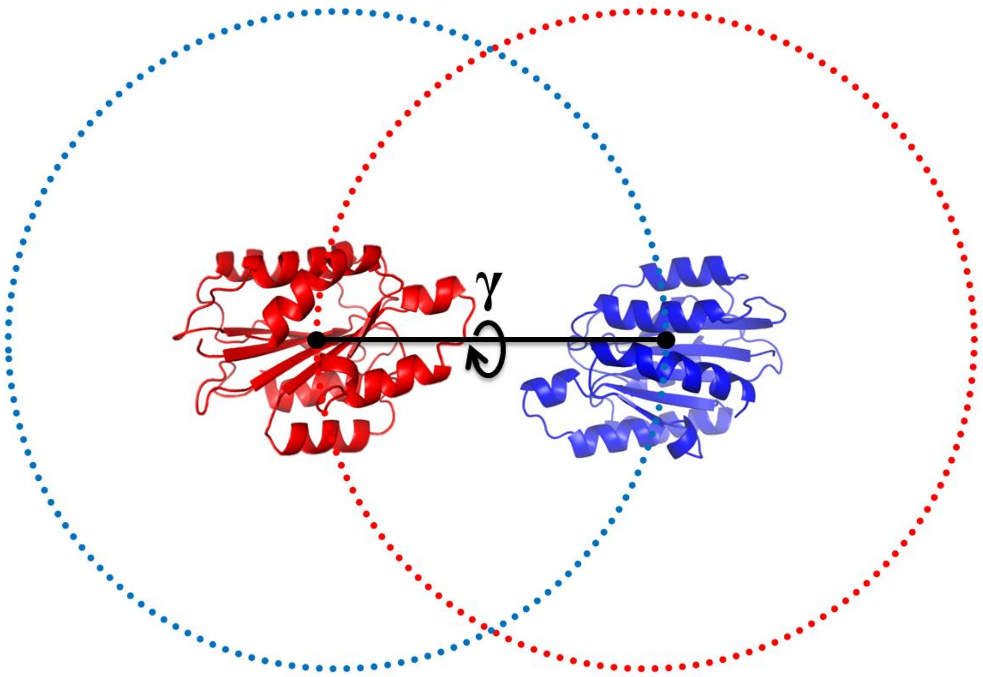


A

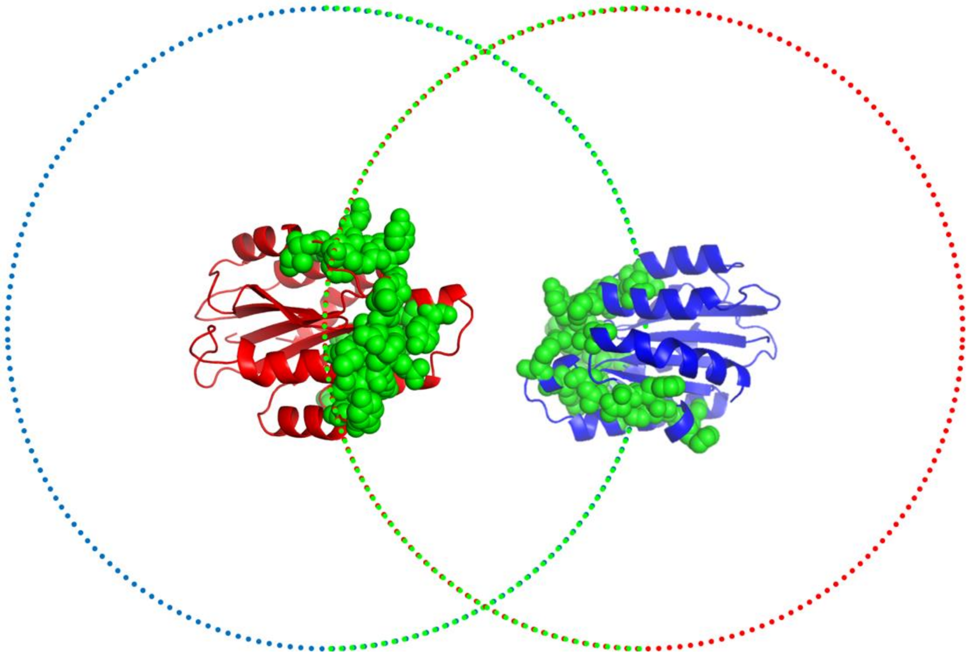

B

**Figure S2:** Starting orientations generation for the rat A1B1 integrin i-domain complex (PDB code 1CK4, 1CK4_A in red and 1CK4_B in blue). The systematically generated starting positions for the receptor and ligand proteins are plotted as blue and red points.

(A) Different starting orientations were generated for each starting positions of the receptor and ligand proteins by applying 5 rotations of the gamma Euler angle defined with the axis connecting the centers of mass of the 2 proteins.

(B) After filtration using JET information, docking calculations are only performed for those starting points that are located in the vicinity of JET predicted interface residues (plotted as green spheres on the proteins), thus considerably reducing the computation time.

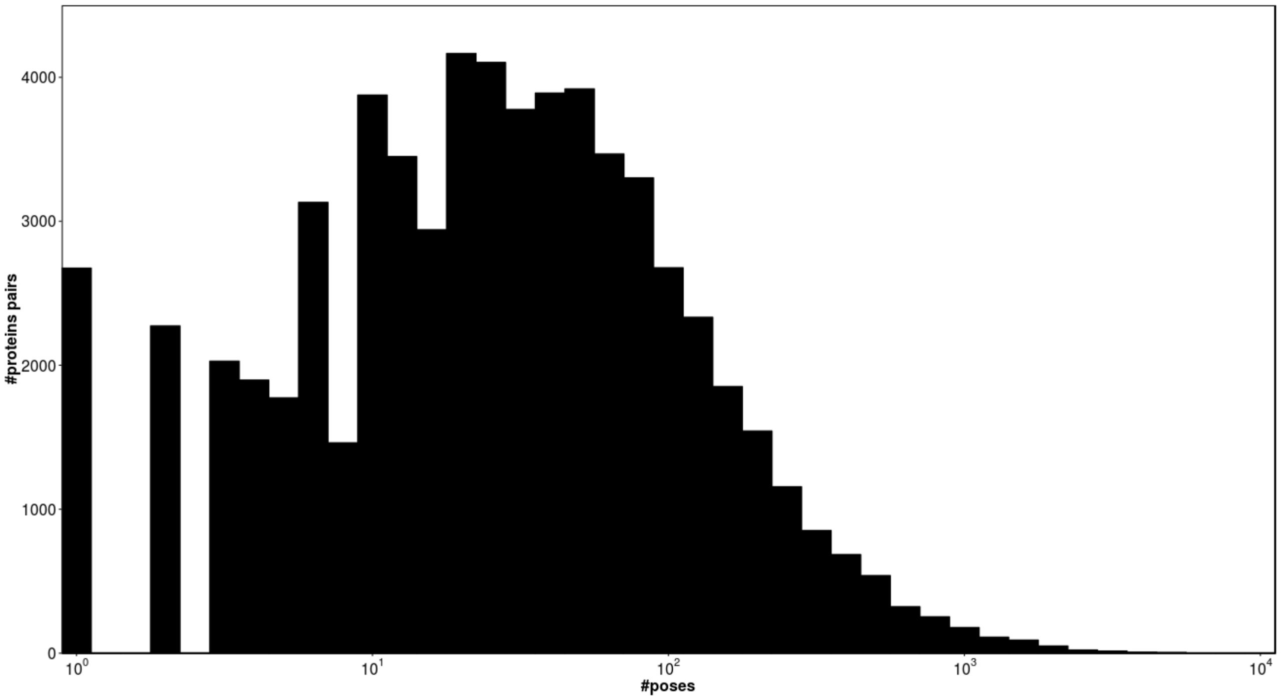


A

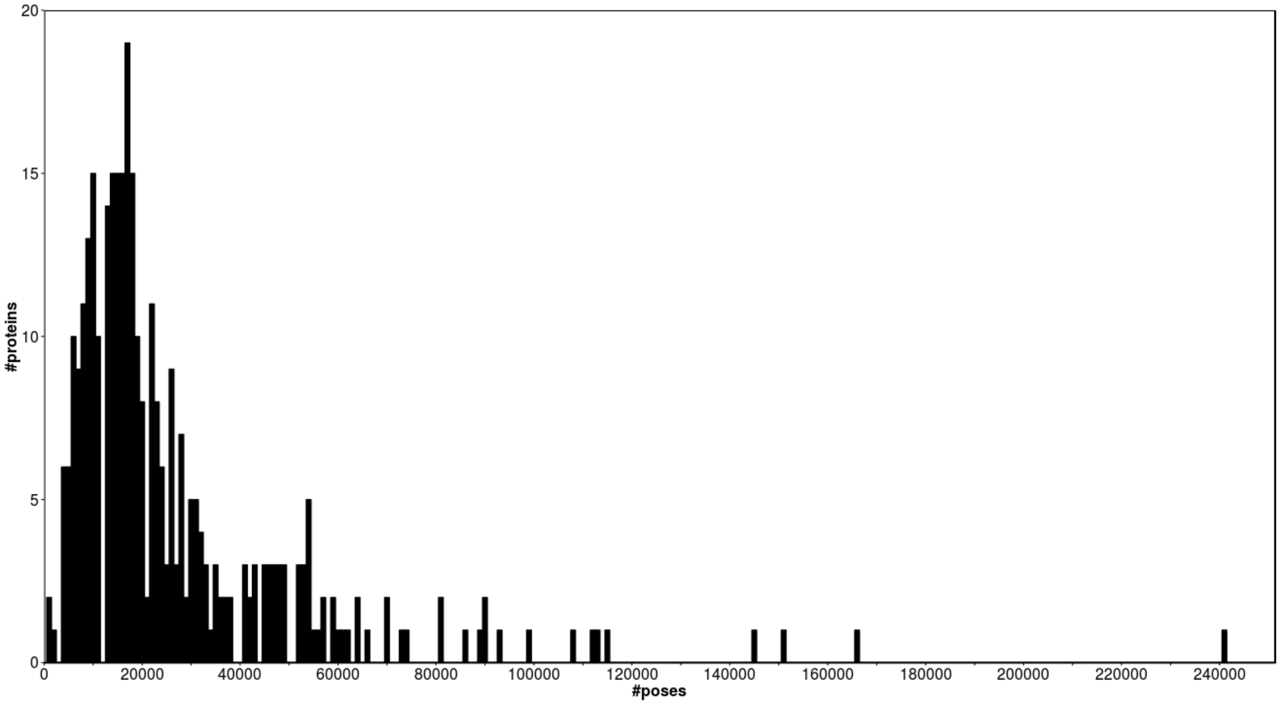


B

**Figure S3:**
(A) Distribution of the number of kept docking poses (using a logarithmic scale) for each protein pair after filtering on the interaction energy.
(B) Final distribution of the number of kept docking poses for each protein (taking into account all its partners) after filtering on the interaction energy.

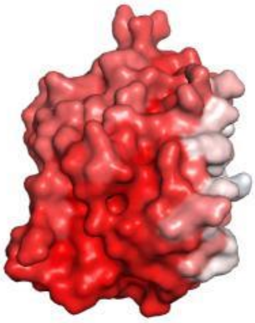

| residue | PIP mean |
|---|---|
| ... | ... |
| 350 | 0.1323255515 |
| 351 | 0.1336589029 |
| 352 | 0.0089175168 |
| 353 | 0 |
| 354 | 0.1346555866 |
| 355 | 0.0951066238 |
| 356 | 0.0082422127 |
| 357 | 0.0760073165 |
| 358 | 0.1340289117 |
| 359 | 0 |
| 360 | 0.0144779084 |
| 361 | 0.1162963671 |
| 362 | 0.1014679365 |
| 363 | 0.0529826201 |
| 364 | 0.0948576183 |
| 365 | 0.111249491 |
| 366 | 0.1071454418 |
| 367 | 0.0989492285 |
| 368 | 0.1022114777 |
| 369 | 0.0620627543 |
| 370 | 0.1023704598 |
| ... | ... |

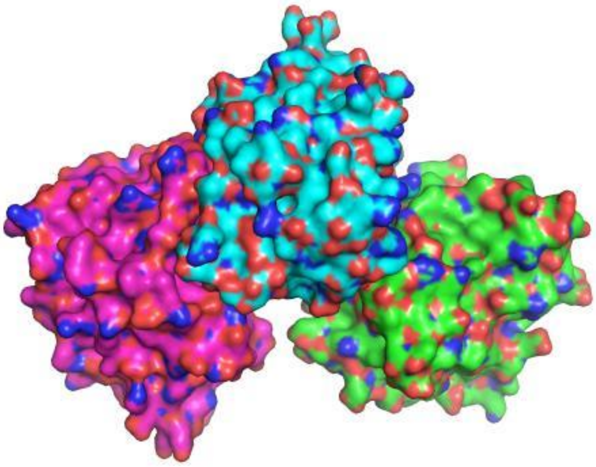

| residue | interface | surface |
|---|---|---|
| 349 | ... | ... |
| 350 | 1 | 1 |
| 351 | 1 | 1 |
| 352 | 0 | 0 |
| 353 | 0 | 0 |
| 354 | 1 | 1 |
| 355 | 1 | 0 |
| 356 | 0 | 0 |
| 357 | 0 | 1 |
| 358 | 1 | 1 |
| 359 | 0 | 0 |
| 360 | 0 | 0 |
| 361 | 0 | 1 |
| 362 | 0 | 1 |
| 363 | 0 | 0 |
| 364 | 0 | 1 |
| 365 | 0 | 1 |
| 366 | 0 | 1 |
| 367 | 0 | 1 |
| 368 | 0 | 1 |
| 369 | 0 | 1 |
| 370 | 0 | 1 |
| ... | ... | ... |

**AUC=0.619**

**Figure S4(A):** *Global interface* (GI) scoring scheme computed for the protein 1IW0_B (represented in cyan in the bottom panel) that presents 2 experimental partners: 1IW0_C and 1IW0_A (respectively in magenta and green in the bottom panel) . The GI score is computed by comparing the PIP values (mapped on the protein's surface in the top panel, high PIP residues showed in white and low PIP residues in red) with one single global reference experimental interface generated by concatening all the existing experimental interfaces. If a residue is part of the experimental interface between the query protein and at least one of the partners, it is tagged as interface residue in the global reference experimental interface (1 in the *interface* column). Conversely, if a residue is not part of any experimental interfaces, it is tagged as non interface residue (0 in the *interface* column).

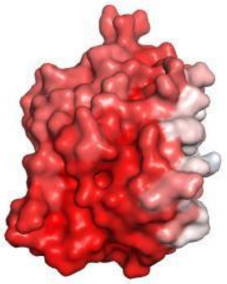

| residue | PIP mean |
|---|---|
| ... | ... |
| 350 | 0.1323255515 |
| 351 | 0.1336589029 |
| 352 | 0.0089175168 |
| 353 | 0 |
| 354 | 0.1346555866 |
| 355 | 0.0951066238 |
| 356 | 0.0082422127 |
| 357 | 0.0760073165 |
| 358 | 0.1340289117 |
| 359 | 0 |
| 360 | 0.0144779084 |
| 361 | 0.1162963671 |
| 362 | 0.1014679365 |
| 363 | 0.0529826201 |
| 364 | 0.0948576183 |
| 365 | 0.111249491 |
| 366 | 0.1071454418 |
| 367 | 0.0989492285 |
| 368 | 0.1022114777 |
| 369 | 0.0620627543 |
| 370 | 0.1023704598 |
| ... | ... |

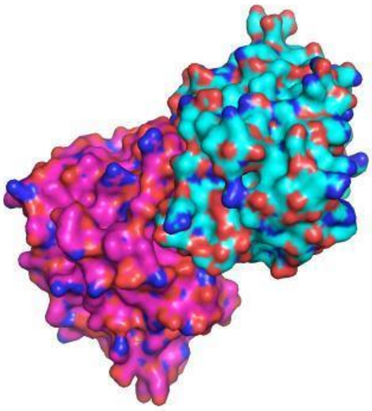

| residue | interface | surface |
|---|---|---|
| ... | ... | ... |
| 350 | 1 | 1 |
| 351 | 1 | 1 |
| 352 | 0 | 0 |
| 353 | 0 | 0 |
| 354 | 1 | 1 |
| 355 | 1 | 0 |
| 356 | 0 | 0 |
| 357 | 0 | 1 |
| 358 | 1 | 1 |
| 359 | 0 | 0 |
| 360 | 0 | 0 |
| 361 | 0 | 1 |
| 362 | 0 | 1 |
| 363 | 0 | 0 |
| 364 | 0 | 1 |
| 365 | 0 | 1 |
| 366 | 0 | 1 |
| 367 | 0 | 1 |
| 368 | 0 | 1 |
| 369 | 0 | 1 |
| 370 | 0 | 1 |
| ... | ... | ... |

AUC=0.759

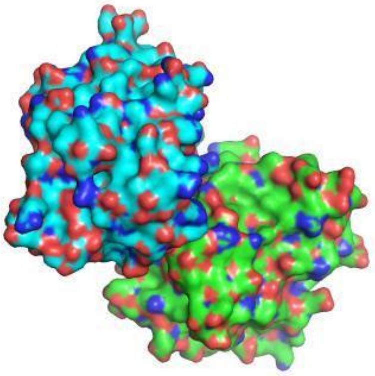

| residue | interface | surface |
|---|---|---|
| ... | ... | ... |
| 350 | 0 | 1 |
| 351 | 0 | 1 |
| 352 | 0 | 0 |
| 353 | 0 | 0 |
| 354 | 0 | 1 |
| 355 | 0 | 0 |
| 356 | 0 | 0 |
| 357 | 0 | 1 |
| 358 | 0 | 1 |
| 359 | 0 | 0 |
| 360 | 0 | 0 |
| 361 | 0 | 1 |
| 362 | 0 | 1 |
| 363 | 0 | 0 |
| 364 | 0 | 1 |
| 365 | 0 | 1 |
| 366 | 0 | 1 |
| 367 | 0 | 1 |
| 368 | 0 | 1 |
| 369 | 0 | 1 |
| 370 | 0 | 1 |
| ... | ... | ... |

AUC=0.516

**Figure S4(B):** *Best interface* (BI) scoring scheme computed for the protein 1IW0_B (represented in cyan in the middle and bottom panels) that presents 2 experimental partners: 1IW0_C (in magenta in the middle panel) and 1IW0_A (in green in the bottom panel). The BI score is computed by comparing the PIP values (mapped on the protein's surface in the top panel, high PIP residues showed in blue and low PIP residues in red) to each reference experimental interface separately, and only the predicted interface associated with the best binding site prediction performance was kept.

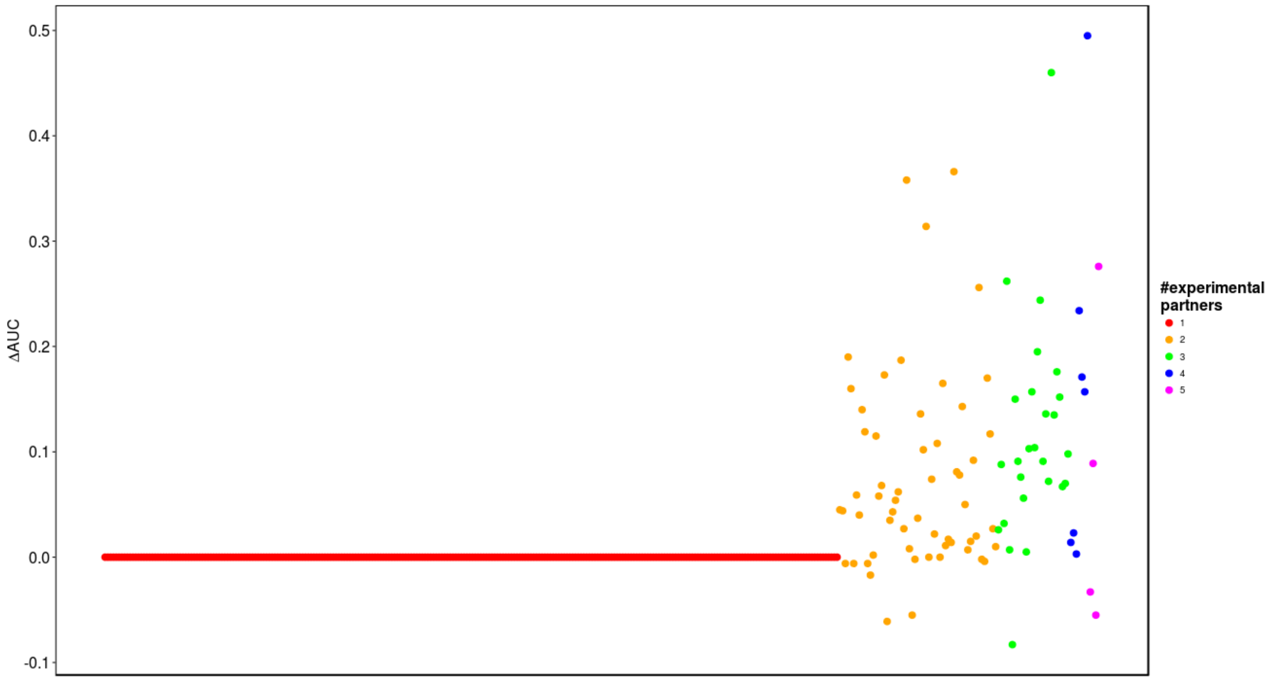


**Figure S5:** Variation of AUC observed for each protein (the x-axis represents the proteins of the SubHCMD dataset ranked according to their number of partners) between the GI score and the BI score ($\Delta AUC = AUC_{BI} - AUC_{GI}$), the dots colors indicate the number of experimental partners included in the CC-D for each protein.

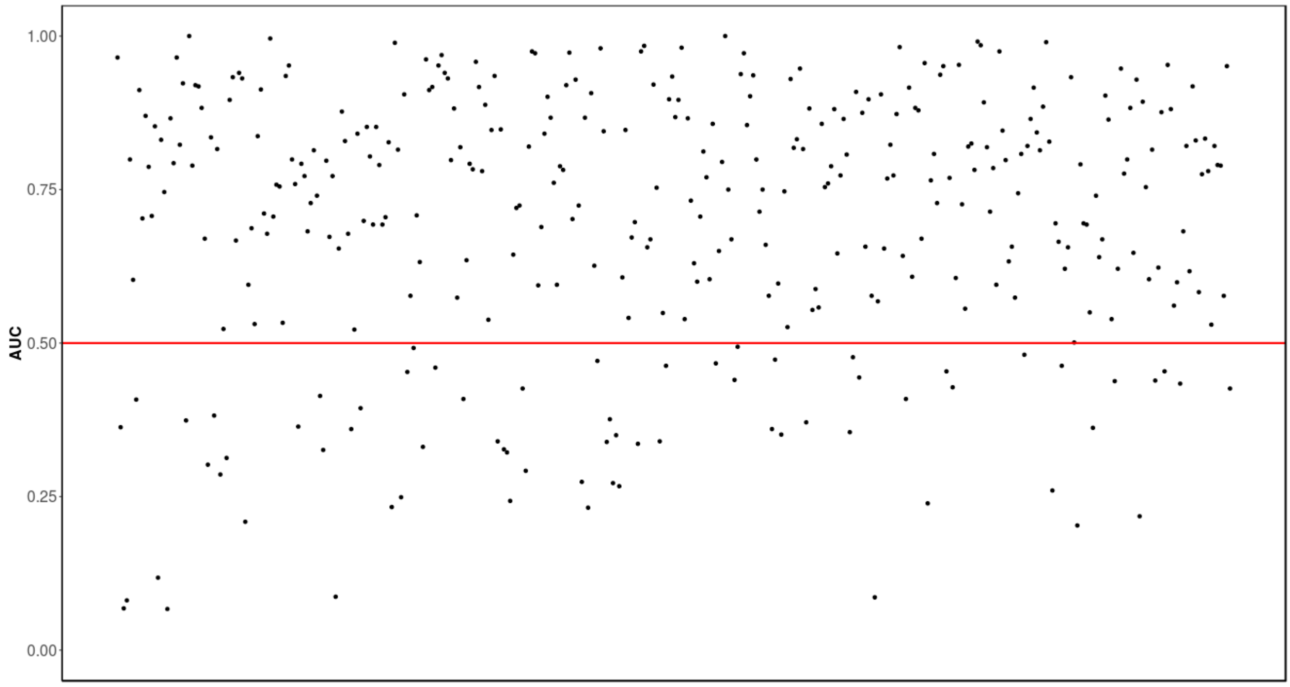


**Figure S6:** AUC value for each protein in our CC-D dataset obtained with the BI score. The red line represents the random threshold of 0.5.

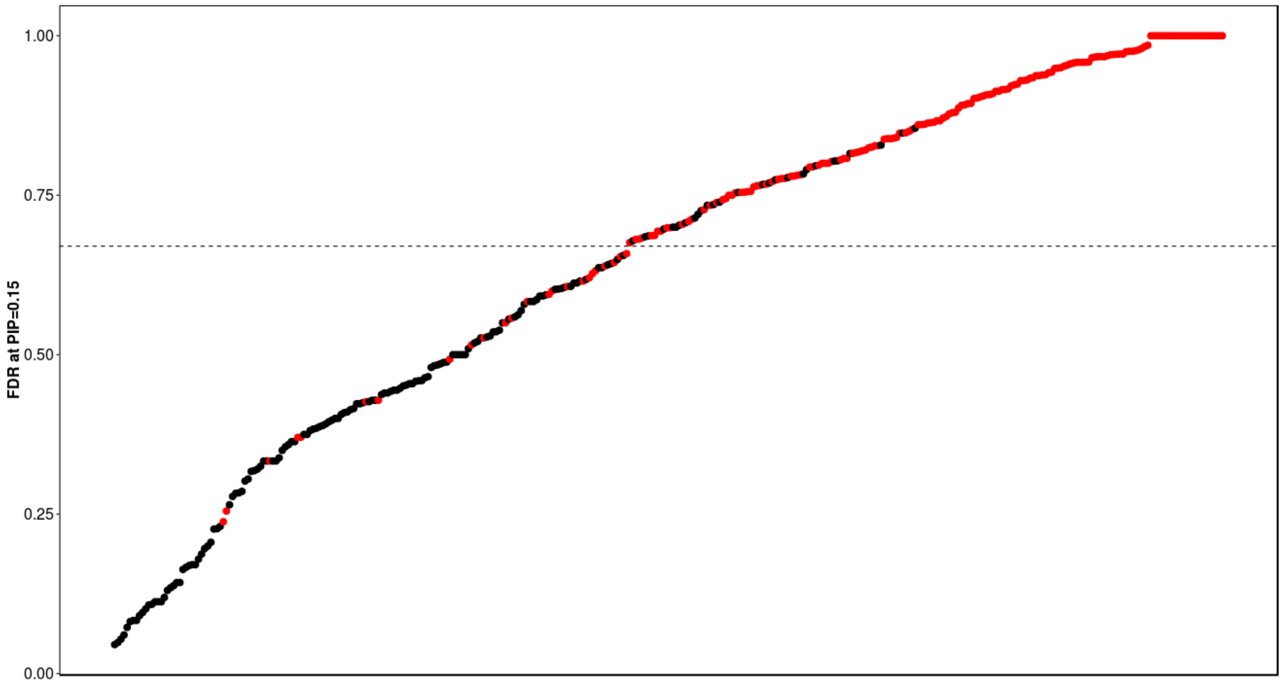


**Figure S7:** Correlation between the visual protein inspection and the proteins FDR (FP/P) values computed at the optimal PIP cut-off of 0.15. The red dots represent proteins for which predicted alternate interfaces were visually observed by mapping the PIP values on the proteins' surface whereas the black dots point out proteins with no observed predicted alternate interface. Proteins were ordered according to their FDR values. The dot line represents the optimal FDR threshold of 0.67.

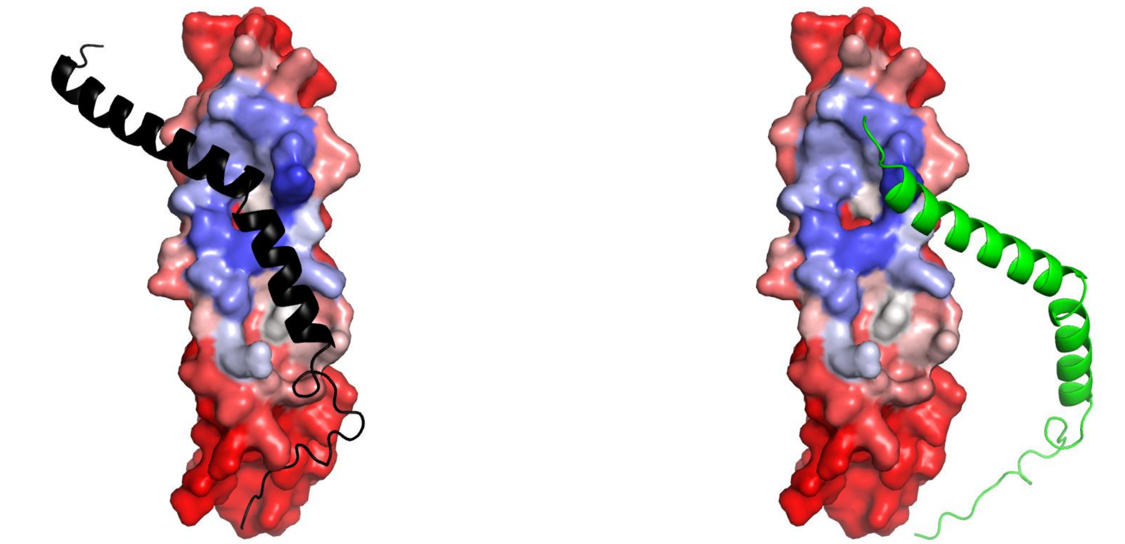

A

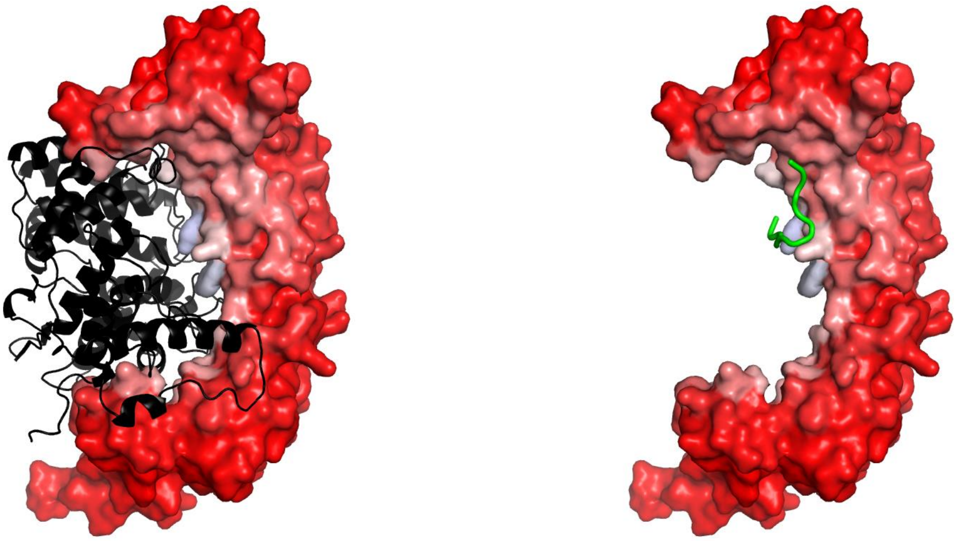

B

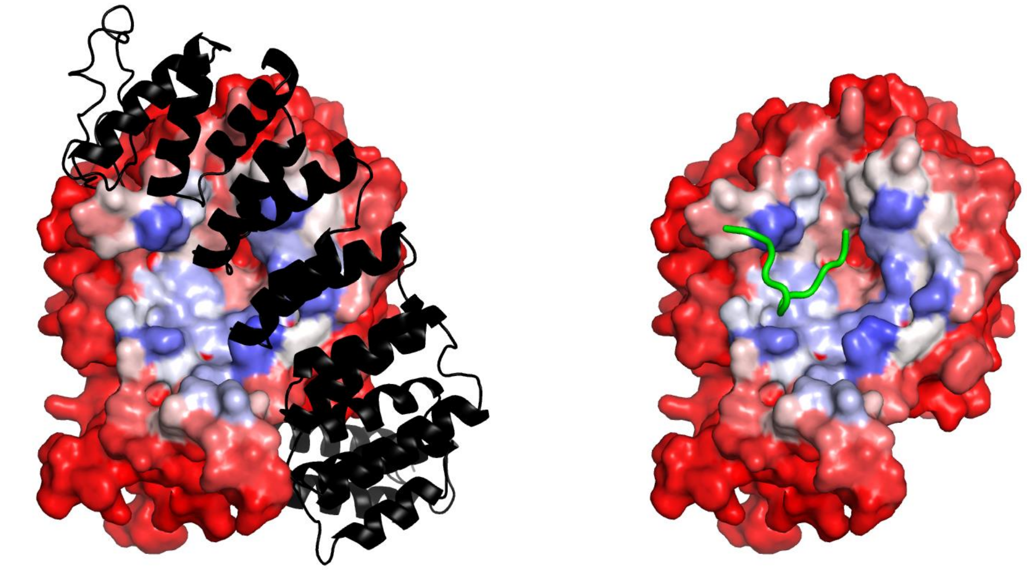

C

**Figure S8** (continued next page)

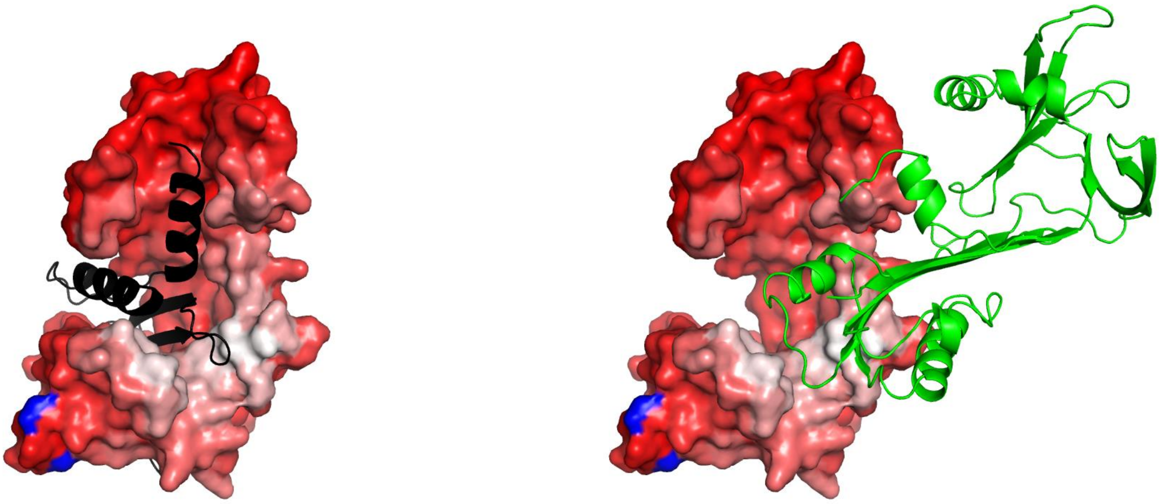

D

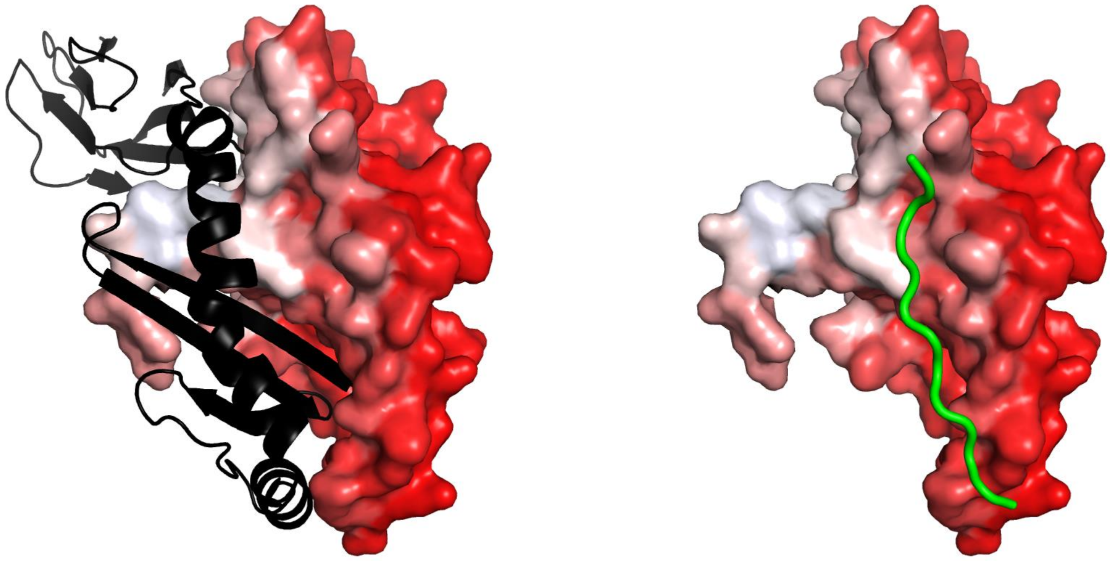

E

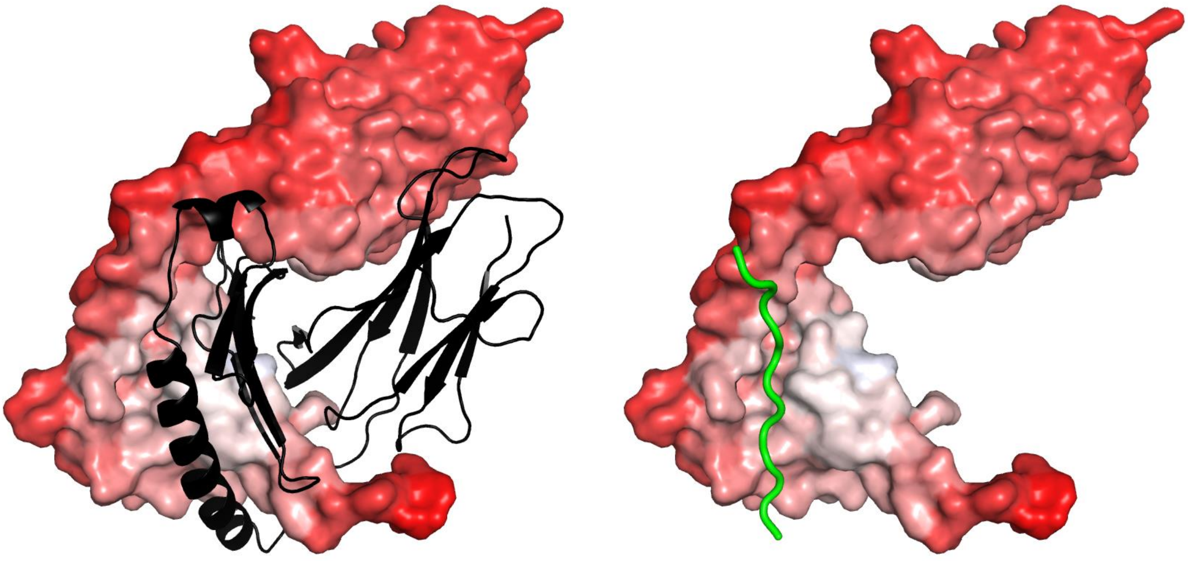

F

**Figure S8** (continued next page)

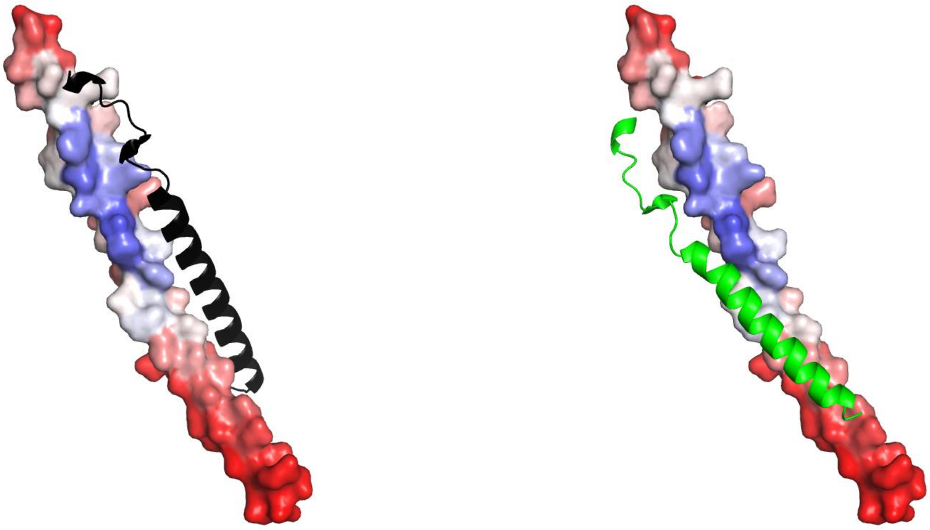

G

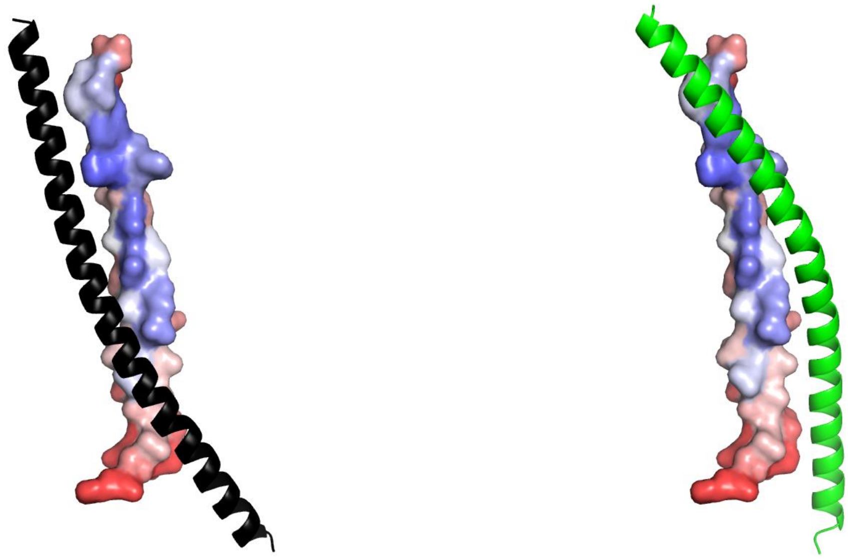

H

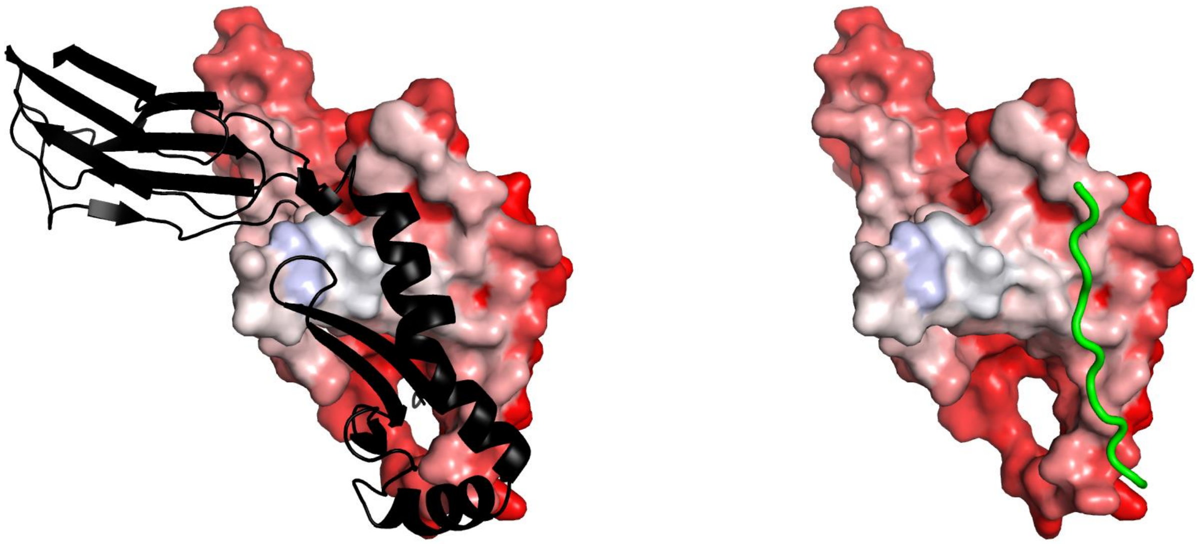

I

**Figure S8** (continued next page)

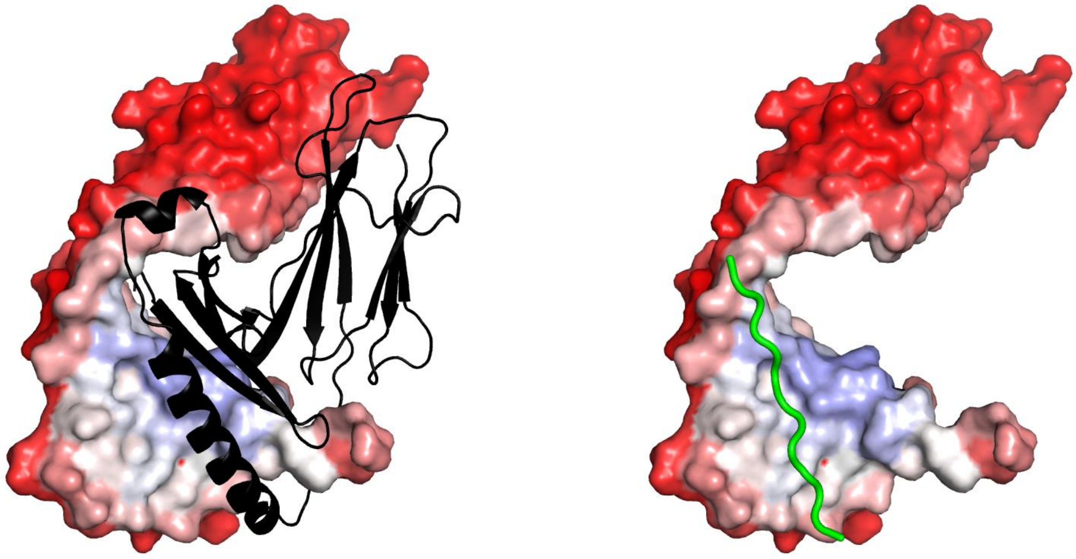

J

**Figure S8(A_J):** Mapping the PIP values on a protein's surface, high PIP residues are shown in blue and low PIP residues are shown in red. The reference experimental partner (ref) is shown in a black cartoon representation, while the alternate partner (proteic or peptidic) (alt) is shown in green.
(A) 1AVO_B with 1AVO_A(ref) and 1AVO_C(alt).
(B) 1D8D_A with 1D8D_B(ref) and 1D8D_P(alt).
(C) 1D8D_B with 1D8D_A(ref) and 1D8D_P(alt).
(D) 1JJO_C with 1JJO_A(ref), 1JJO_E(ref) and 1JJO_D(alt).
(E) 2NNA_A with 2NNA_B(ref) and 2NNA_C(alt).
(F) 2NNA_B with 2NNA_A(ref) and 2NNA_C(alt).
(G) 3BRT_B with 3BRT_C(ref) and 3BRT_A(alt).
(H) 3BRT_C with 3BRT_B(ref) and 3BRT_D(alt).
(I) 3C5J_A with 3C5J_B(ref) and 3C5J_C(alt).
(J) 3C5J_B with 3C5J_A(ref) and 3C5J_C(alt)

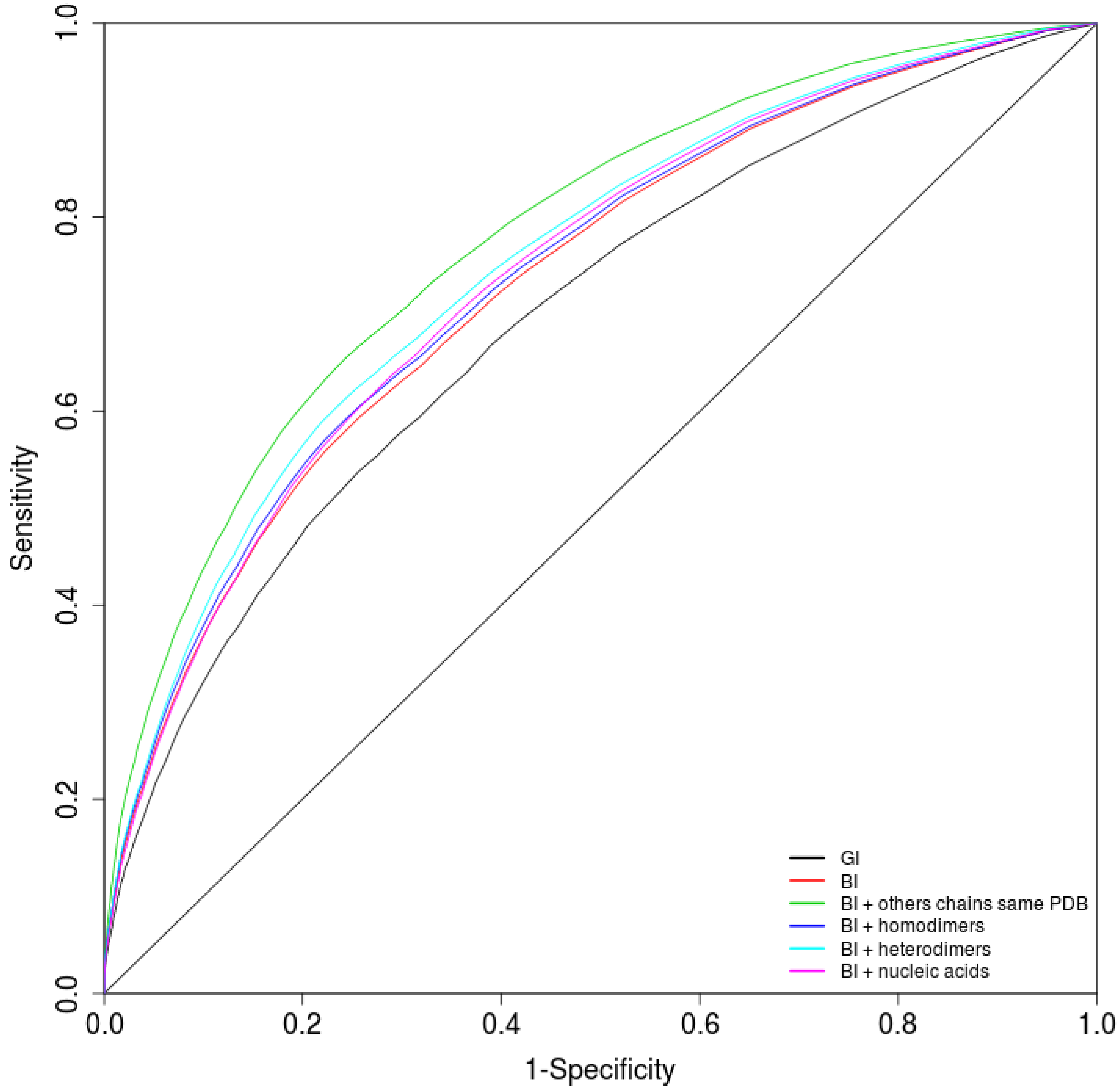


**Figure S9:** ROC curves of the PIP prediction obtained with the GI score with the reference experimental interfaces (REI) (black line), with the BI score with the REI (red line), with the BI score with REI and alternate experimental interfaces from other chains of the same PDB (green line), with the BI score with REI and alternate experimental interfaces from homodimers (blue line), with the BI score with REI and alternate experimental interfaces from heteromdimers (cyan line) and with the BI score with REI and alternate nucleic acid experimental interfaces (magenta line). The diagonal dotted line corresponds to random prediction.

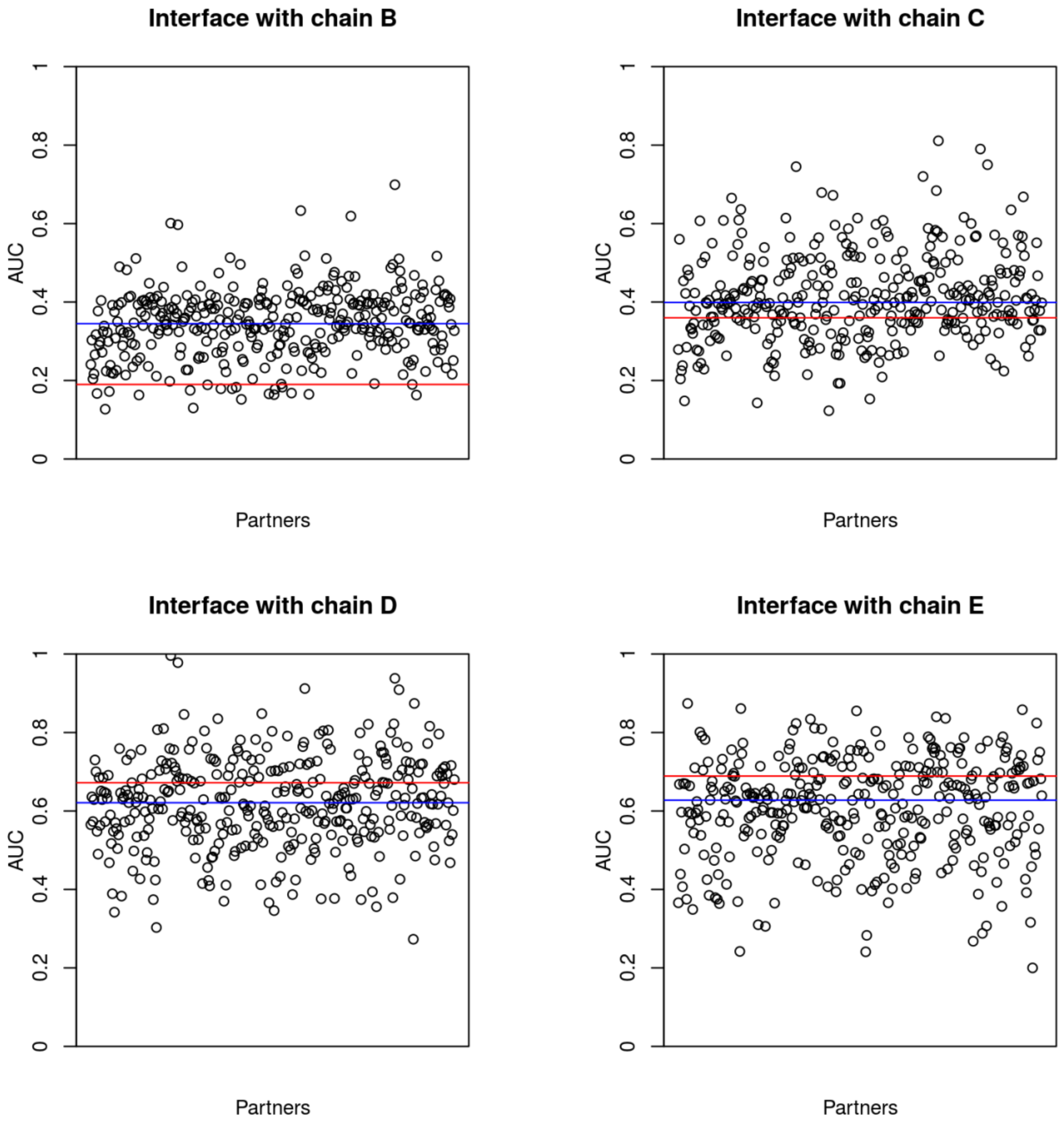


A

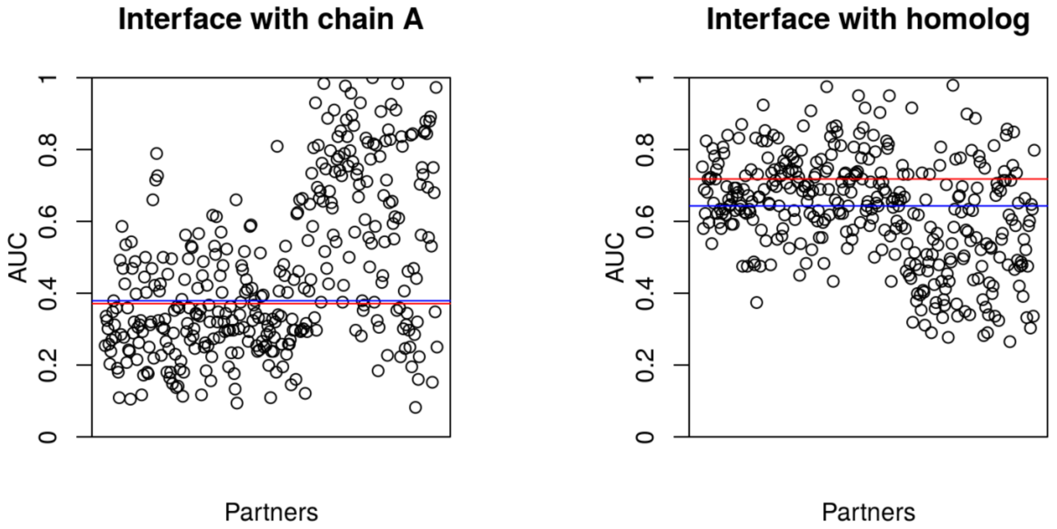


B

**Figure S10** (continued next page)

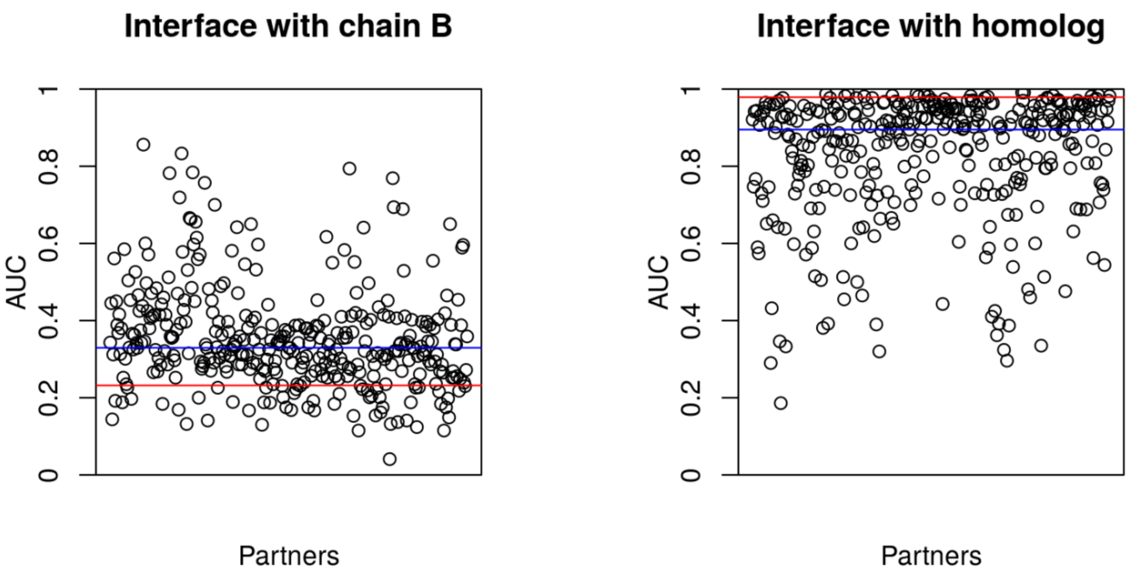


C

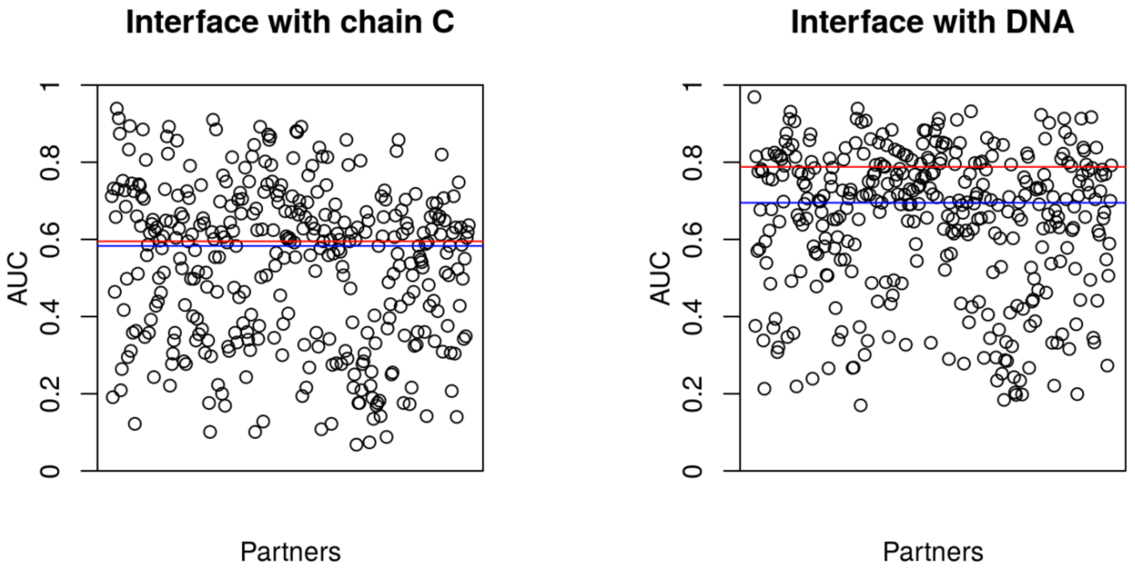


D

**Figure S10**: Individual AUC values obtained by single docking for proteins (A) 1LI1_F, (B) 2P1L_B, (C) 2BOV_A and (D) 1GT0_D for their different experimental interfaces. The red line represents the value of the global AUC computed with the BI score on the cross-docking results and the blue line represents the median value of the individual AUC obtained by single docking.

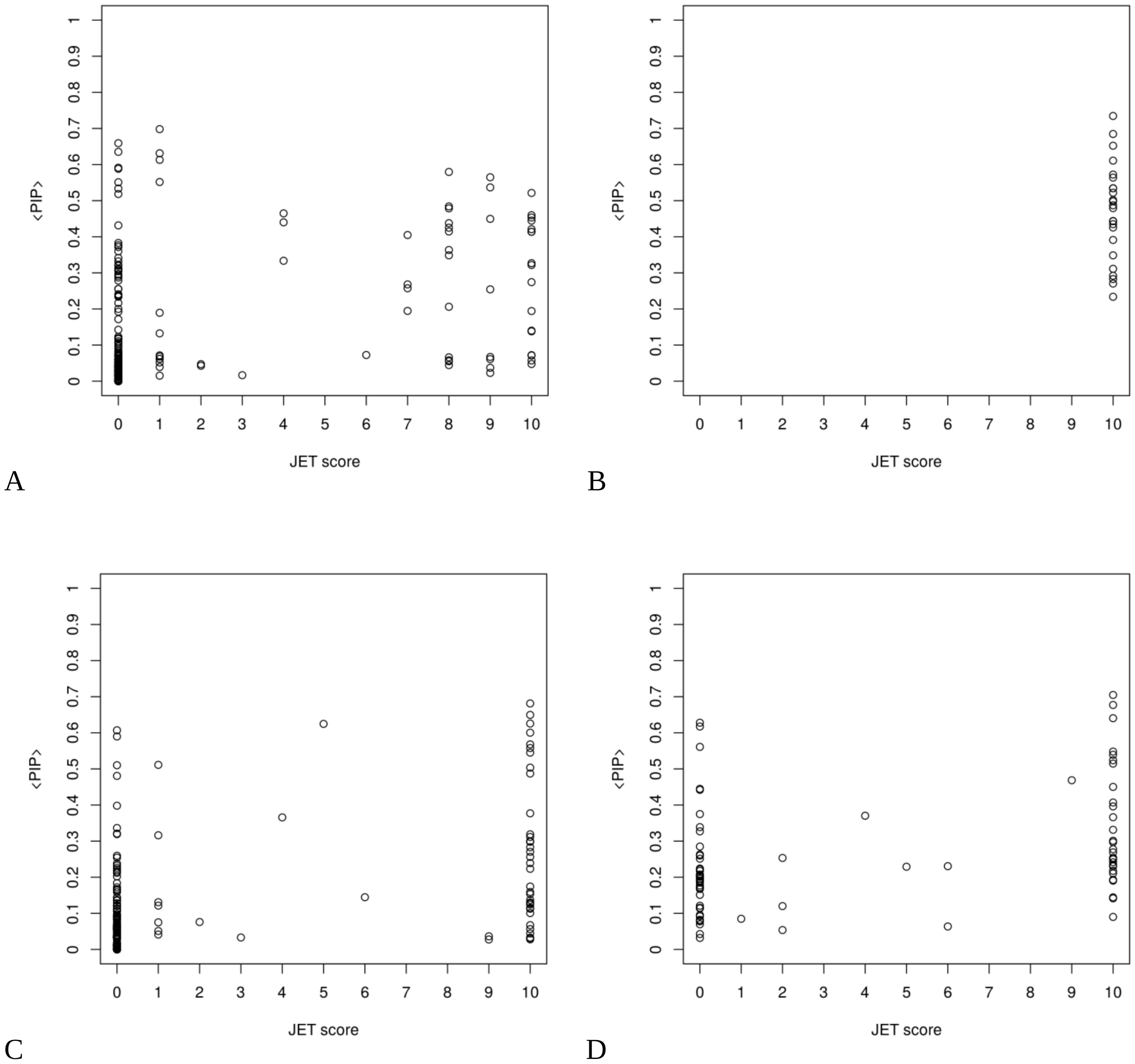


**Figure S11**: Distribution of the <PIP> values as a function of the JET score for proteins (A) 1LI1_F, (B) 2P1L_B, (C) 2BOV_A and (D) 1GT0_D. It is to note that the 2P1L_B protein is a very small protein (24 residues) which may explain the specific profile observed for this protein (all residues obtained a JET score of 10).

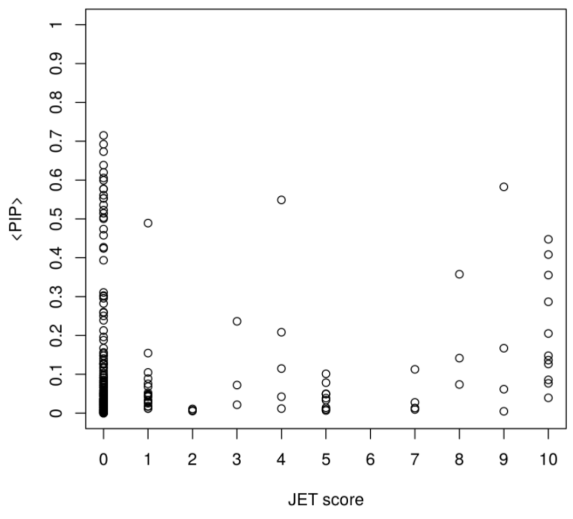


A

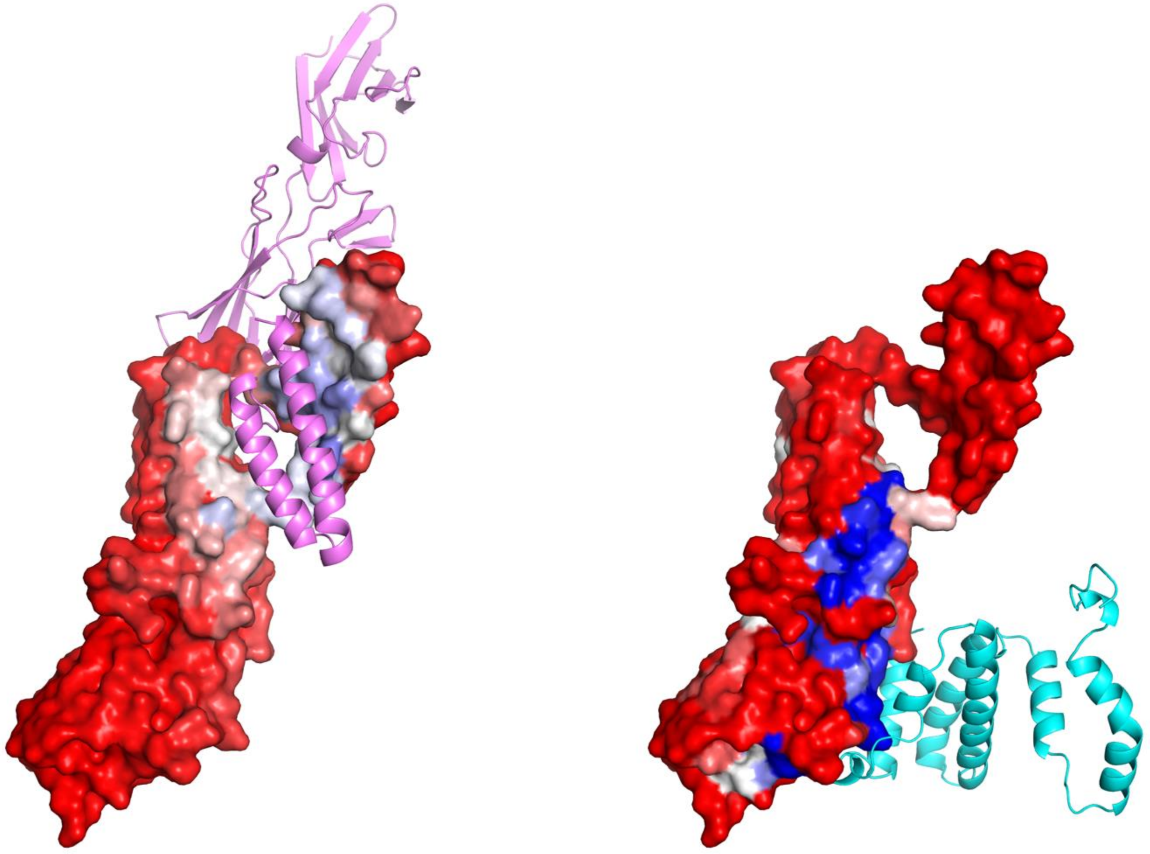

B

**Figure S12: (A)** Distribution of the <PIP> values as a function of the JET score for protein 3BES_R. (B) On the left panel, <PIP> values are mapped on the protein's surface (high <PIP> residues are shown in blue and low <PIP> residues are shown in red) and 3BES_D is showed in cartoon representation in violet. On the right panel, JET score values are mapped on the protein's surface (high JET score residues are shown in blue and low JET score residues are shown in red) and 3BES_L is showed in cartoon representation in cyan.

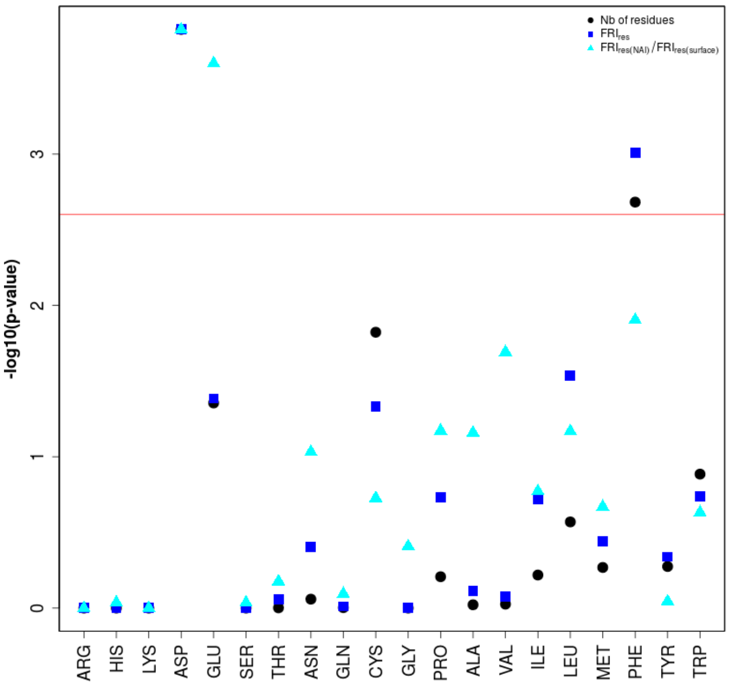


A

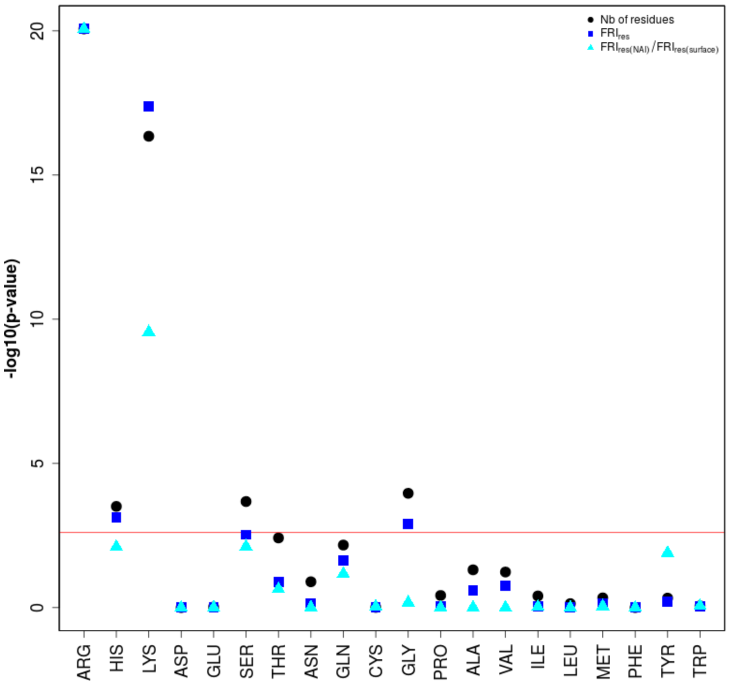


B

**Figure S13:** Residues are plotted as the distribution of the negative logarithmic p-value (-log10(p-value)) of the one-tailed Wilcoxon test for the number of each residue type in the interface (black dot), for the fraction of each residue type in the interface (FRIres(type)) (blue square) and for the fraction of each residue type in the interface ($FRI_{res(type)}$) in the NAI compared to the fraction of each residue type observed in the whole corresponding protein surface ($FRI_{res(type)[surface]}$) (cyan triangle) (A) option « less », (B) option « greater ». All points located above the Bonferroni threshold (red line) display a mean value significantly inferior (A) or superior (B) in the NAI than in the PPI.

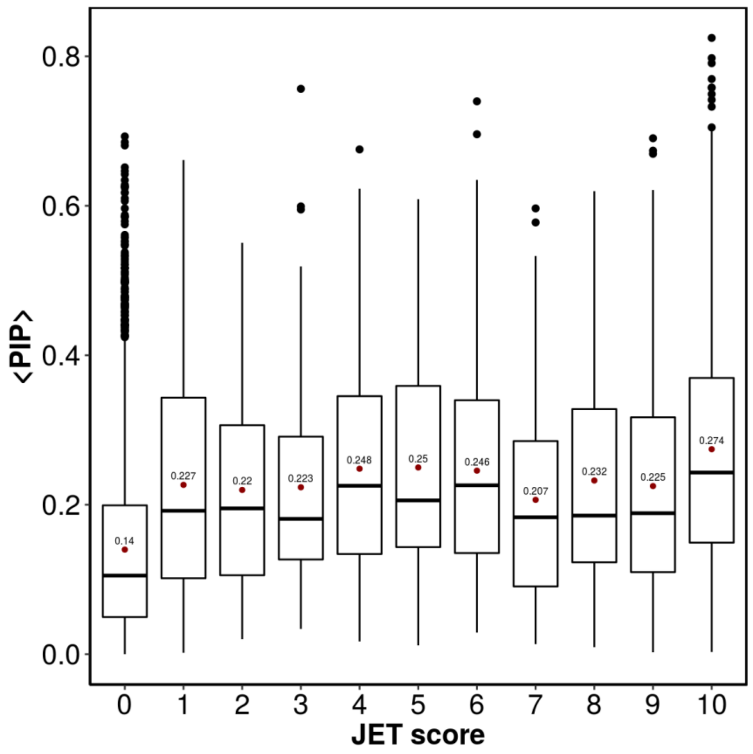


**Figure S14:** Boxplot of the distribution of <PIP> values as a function of JET score for proteins of the SubHCMD dataset with a nucleic acid binding site (1GT0_C, 1GT0_D, 1MDM_A, 1MDM_B, 1WSU_A, 1WSU_B, 1WSU_C, 2NQB_D, 2NQB_G, 2QOU_C, 2QOU_D, 2QOU_E, 2QOU_F, 2QOU_G, 2QOU_H, 2QOU_I, 2QOU_J, 2QOU_K, 2QOU_L, 2QOU_M, 2QOU_N, 2QOU_O, 2QOU_Q, 1QOU_R, 2QOU_S, 2QOU_U, 2QOV_1, 2QOV_3, 2QOV_4, 2QOV_D, 2QOV_E, 2QOV_G, 2QOV_H, 2QOV_K, 2QOV_L, 2QOV_M, 2QOV_N, 2QOV_P, 2QOV_T, 2QOV_V, 2QOV_X, 2QOV_Z, 3B6F_C, 3B6F_H). The red dot and its corresponding label indicates the mean value of <PIP> for each group of JET score.

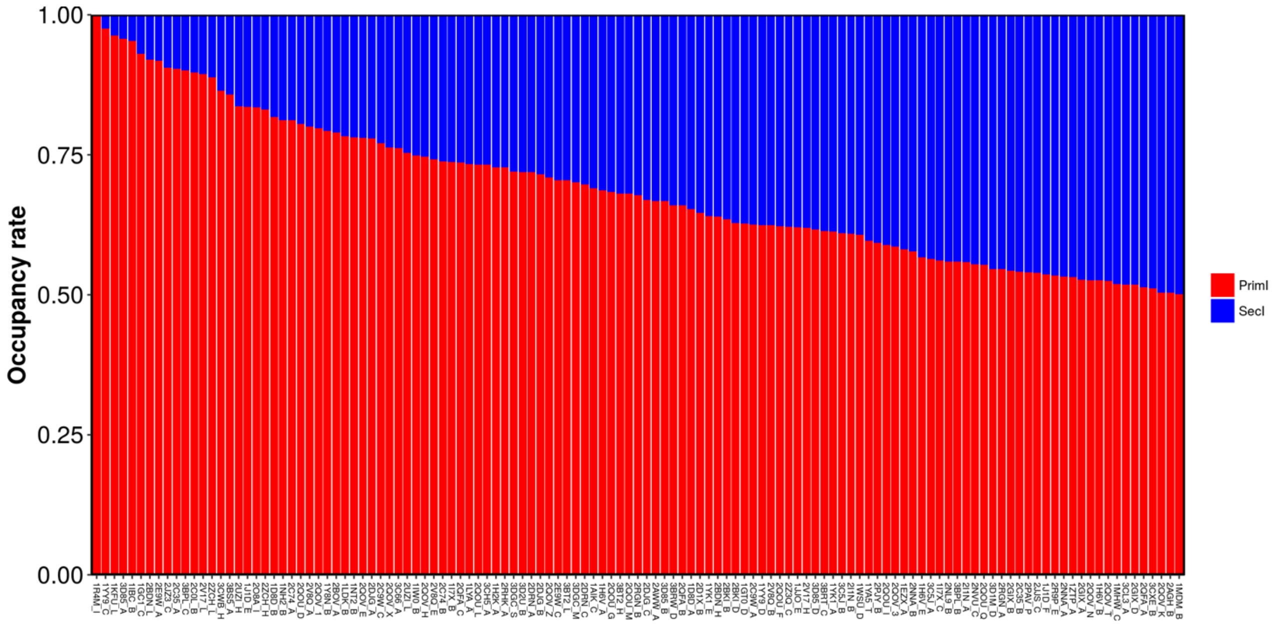


A

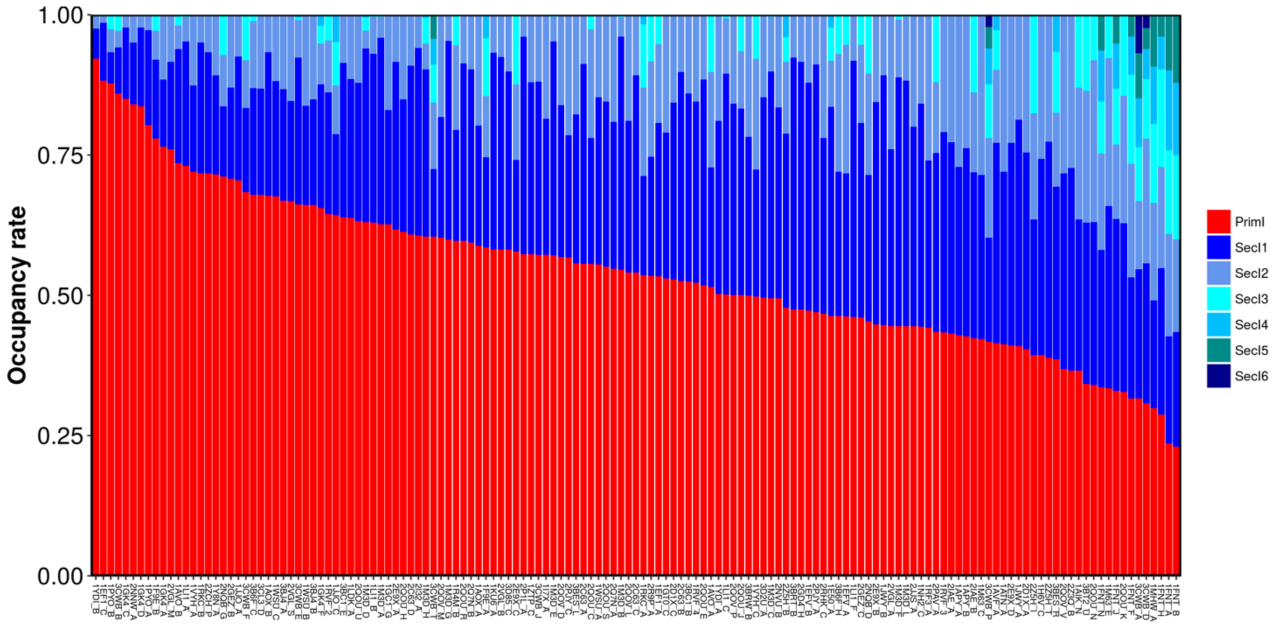


B

**Figure S15**: Occupancy rates for proteins with 2 interfaces (A) or more (B) computed for each interface of each protein as the number of docking poses that presents the best AUC value for this binding site divided by the total number of poses.

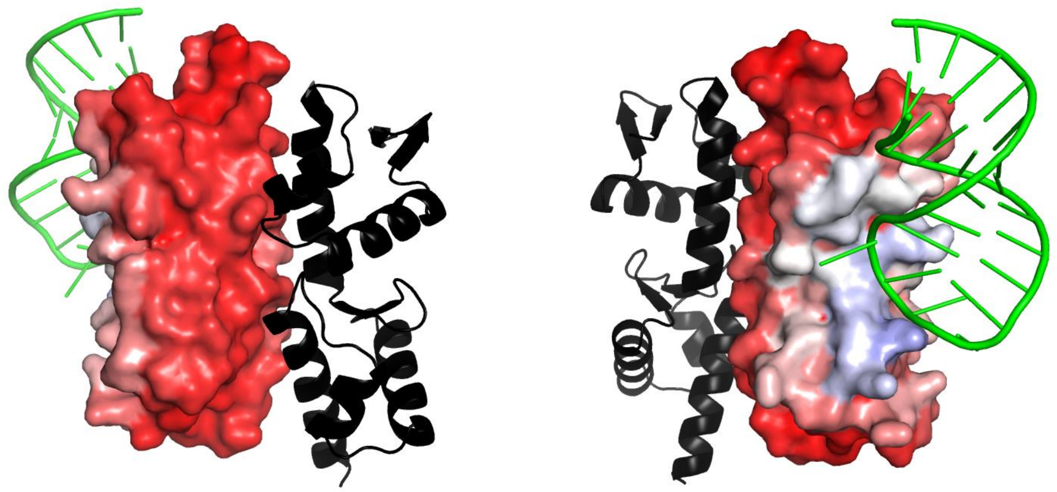

A

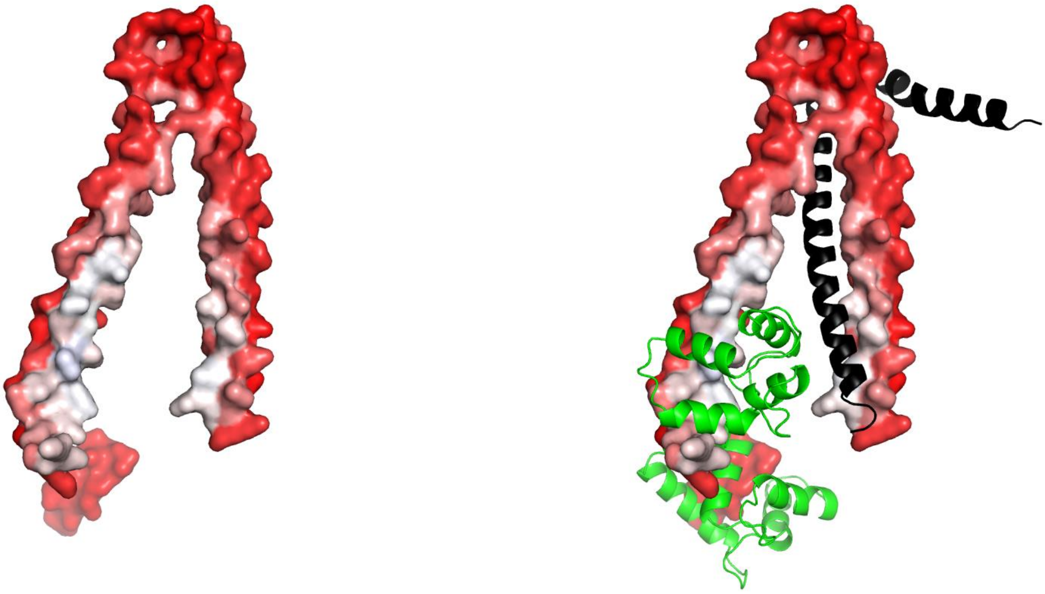

B

**Figure S16:** Proteins with multiple predicted interfaces can present
(A) one binding site predominantly targeted by the different partners during the cross-docking simulations or
(B) similar occupancy rates observed between the different predicted binding sites.
The PIP values are mapped on the protein's surface (high PIP residues are shown in blue and low PIP residues are shown in red) and the different partners are showed in cartoon representation in green and black.

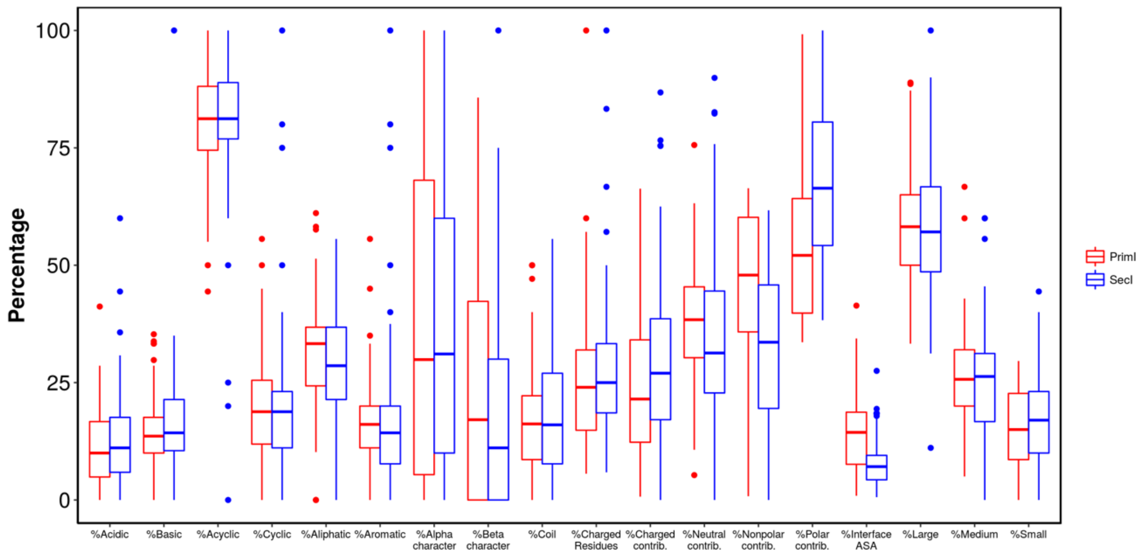


A

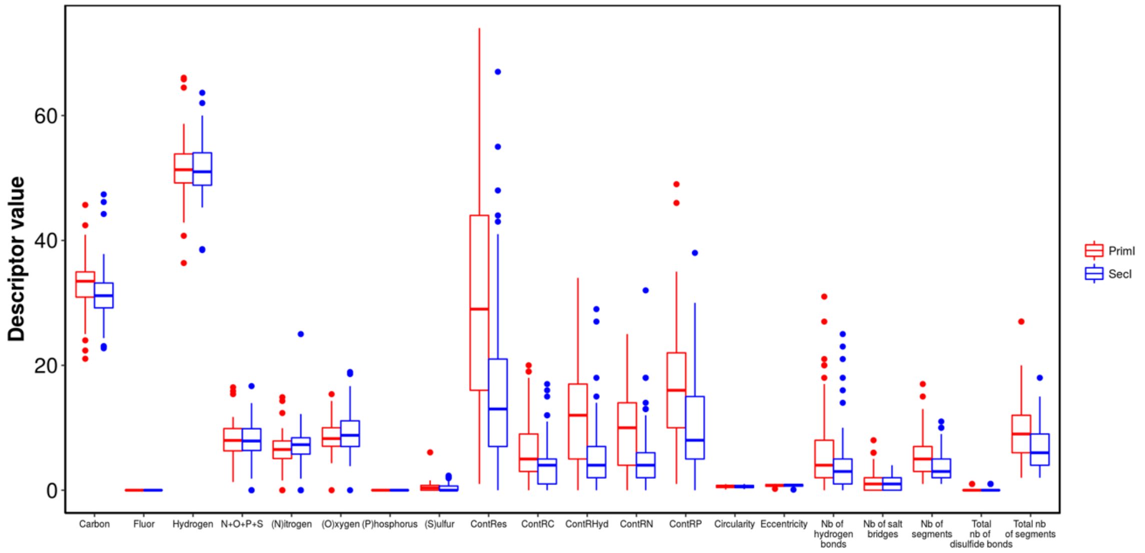


B

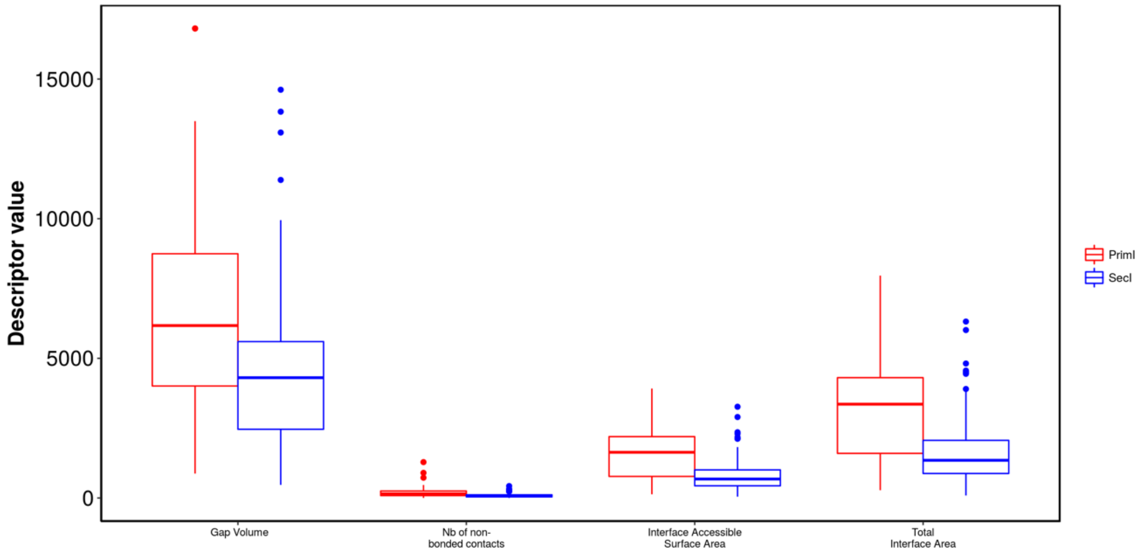


C

**Figure S17(A-C):** Boxplot of the distribution of the values obtained for each of the 43 P2PI inspector descriptors with the 85 primary interface (PrimI) (showed in red) and the 85 secondary interface (SecI) (showed in blue).

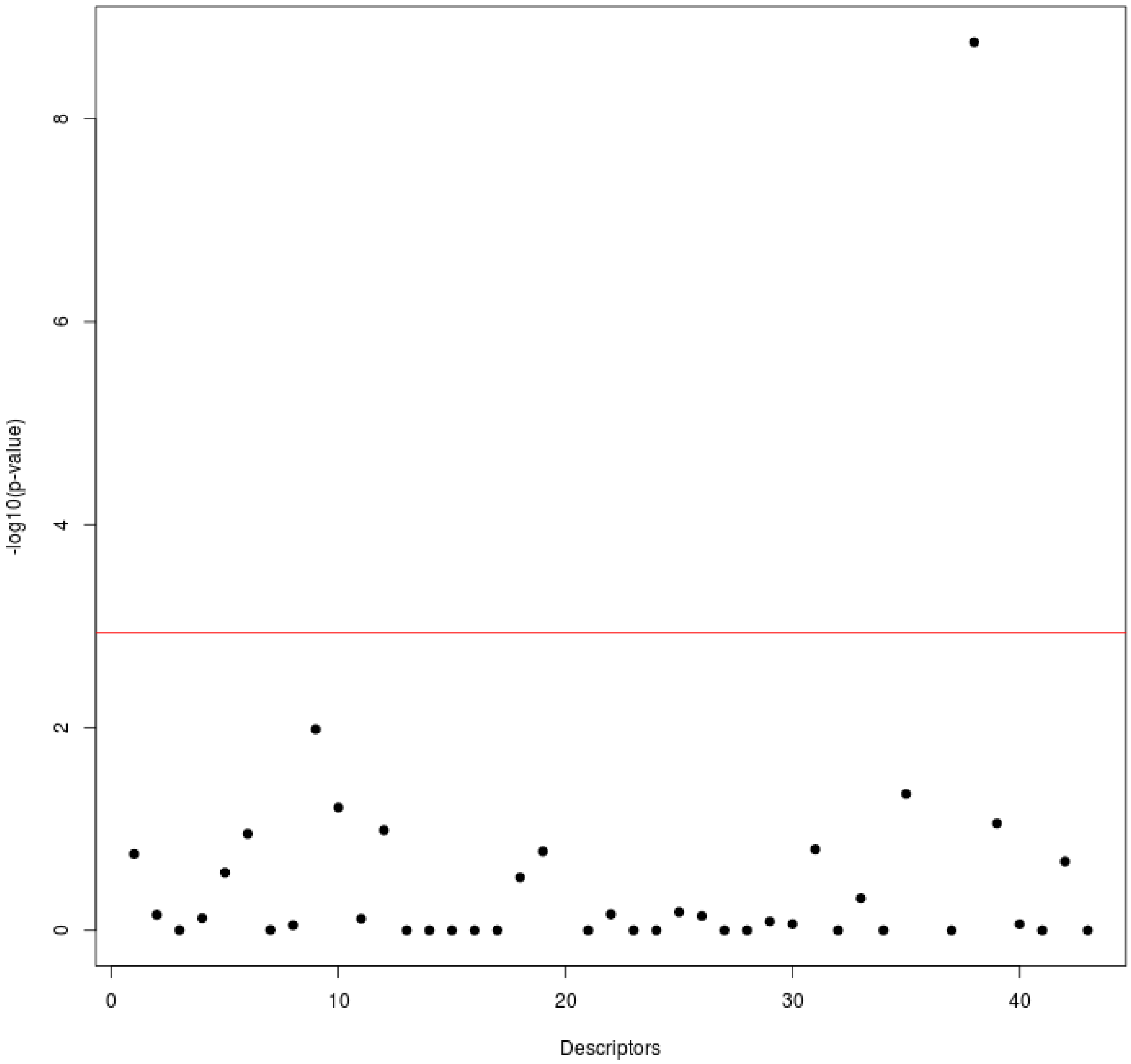


**Figure S18:** The 43 P2PI inspector descriptors are plotted as the distribution of the negative logarithmic p-value (-log10(p-value)) of the one-tailed Student test option « less ». All descriptors located above the Bonferroni threshold (red line) display a mean value significantly inferior in the PrimI than in the SecI.

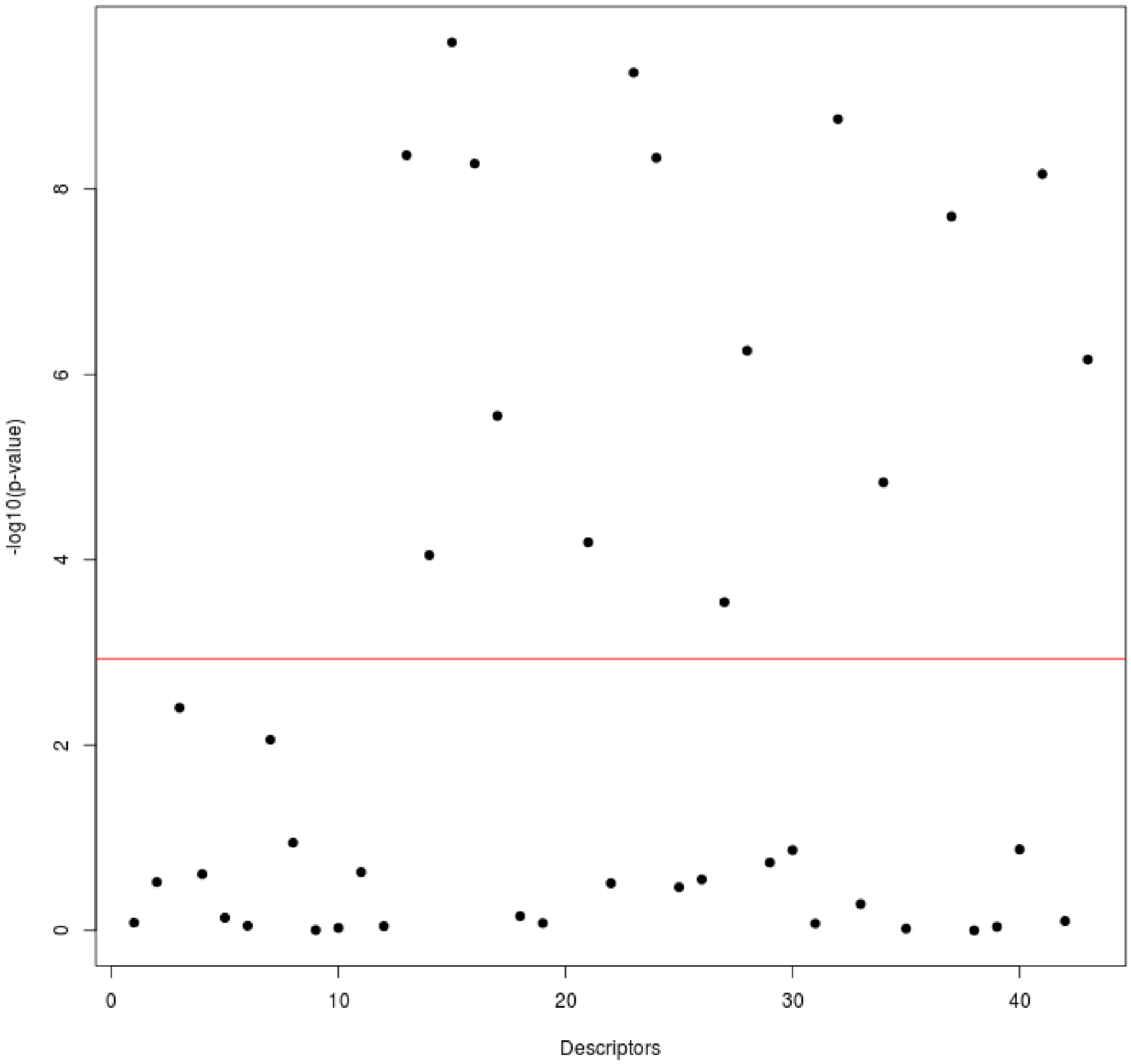


**Figure S19:** The 43 P2PI inspector descriptors are plotted as the distribution of the negative logarithmic p-value (-log10(p-value)) of the one-tailed Student test option « greater ». All descriptors located above the Bonferroni threshold (red line) display a mean value significantly superior in the PrimI than in the SecI.

| PDB | Resolution | Chains | Nb residues | Couples |
|---|---|---|---|---|
| 1AIK | 2.0 | C<br>N | 34<br>36 | 1AIK_C--1AIK_N |
| 1AOX | 2.1 | A<br>B | 201<br>201 | 1AOX_A--1AOX_B |
| 1APY | 2.0 | A<br>B | 161<br>141 | 1APY_A--1APY_B |
| 1ATN | 2.8 | A<br>D | 372<br>258 | 1ATN_A--1ATN_D |
| 1AVF | 2.36 | A<br>P | 322<br>21 | 1AVF_A--1AVF_P |
| 1AVO | 2.8 | A<br>B | 60<br>140 | 1AVO_A--1AVO_B |
| 1CK4 | 2.2 | A<br>B | 193<br>195 | 1CK4_A--1CK4_B |
| 1D8D | 2.0 | A<br>B | 323<br>407 | 1D8D_A--1D8D_B |
| 1E50 | 2.6 | A<br>D | 115<br>125 | 1E50_A--1E50_D |
| 1EF1 | 1.9 | A<br>C | 289<br>87 | 1EF1_A--1EF1_C |
| 1EFV | 2.1 | A<br>B | 312<br>252 | 1EFV_A--1EFV_B |
| 1EZX | 2.6 | A<br>B<br>C | 335<br>36<br>140 | 1EZX_A--1EZX_B<br>1EZX_A--1EZX_C |
| 1F9E | 2.9 | A<br>B | 153<br>89 | 1F9E_A--1F9E_B |
| 1FNT | 3.2 | A<br>B<br>F<br>H<br>J<br>N | 238<br>247<br>233<br>196<br>204<br>233 | 1FNT_A--1FNT_B<br>1FNT_A--1FNT_H<br>1FNT_B--1FNT_J<br>1FNT_F--1FNT_N<br>1FNT_H--1FNT_N |
| 1GC1 | 2.5 | C<br>G | 181<br>297 | 1GC1_C--1GC1_G |
| 1GK4 | 2.3 | A<br>C<br>D<br>F | 79<br>70<br>78<br>74 | 1GK4_A--1GK4_C<br>1GK4_A--1GK4_D<br>1GK4_A--1GK4_F<br>1GK4_C--1GK4_D<br>1GK4_C--1GK4_F<br>1GK4_D--1GK4_F |
| 1GL4 | 2.0 | A<br>B | 273<br>89 | 1GL4_A--1GL4_B |
| 1GT0 | 2.6 | C<br>D | 138<br>80 | 1GT0_C--1GT0_D |

**Table S1.** (continued next page)

| PDB | Resolution | Chains | Nb residues | Couples |
|---|---|---|---|---|
| 1H2K | 2.15 | A | 332 | 1H2K_A--1H2K_S |
| | | S | 24 | |
| 1H6V | 3.0 | A | 490 | 1H6V_A--1H6V_B |
| | | B | 487 | 1H6V_A--1H6V_C |
| | | C | 482 | 1H6V_C--1H6V_E |
| | | E | 491 | |
| 1H8B | NMR | A | 73 | 1H8B_A--1H8B_B |
| | | B | 23 | |
| 1I7X | 3.0 | A | 522 | 1I7X_A--1I7X_B |
| | | B | 57 | 1I7X_A--1I7X_C |
| | | C | 521 | |
| 1IBC | 2.73 | A | 167 | 1IBC_A--1IBC_B |
| | | B | 88 | |
| 1IW0 | 1.4 | A | 207 | 1IW0_A--1IW0_B |
| | | B | 209 | 1IW0_B--1IW0_C |
| | | C | 207 | |
| 1J1D | 2.61 | E | 75 | 1J1D_E--1J1D_F |
| | | F | 116 | |
| 1JJO | 3.06 | A | 40 | 1JJO_A--1JJO_C |
| | | C | 244 | 1JJO_A--1JJO_E |
| | | E | 33 | 1JJO_C--1JJO_E |
| 1JWY | 2.3 | A | 753 | 1JWY_A--1JWY_B |
| | | B | 316 | |
| 1KFU | 2.5 | L | 699 | 1KFU_L--1KFU_S |
| | | S | 184 | |
| 1KU6 | 2.5 | A | 535 | 1KU6_A--1KU6_B |
| | | B | 61 | |
| 1LDK | 3.1 | A | 358 | 1LDK_A--1LDK_B |
| | | B | 366 | |
| 1LI1 | 1.9 | A | 228 | 1LI1_A--1LI1_B |
| | | B | 227 | 1LI1_A--1LI1_C |
| | | C | 224 | 1LI1_B--1LI1_C |
| | | F | 225 | 1LI1_B--1LI1_F |
| | | | | 1LI1_C--1LI1_F |
| 1LM5 | 1.8 | A | 189 | 1LM5_A--1LM5_B |
| | | B | 193 | |
| 1LYA | 2.5 | A | 97 | 1LYA_A--1LYA_B |
| | | B | 241 | |

**Table S1.** (continued next page)

| PDB | Resolution | Chains | Nb residues | Couples |
|---|---|---|---|---|
| 1M3D | 2.0 | A | 223 | 1M3D_A--1M3D_B |
| | | | | 1M3D_A--1M3D_C |
| | | B | 224 | 1M3D_A--1M3D_D |
| | | | | 1M3D_A--1M3D_E |
| | | C | 222 | 1M3D_B--1M3D_C |
| | | D | 225 | 1M3D_B--1M3D_D |
| | | | | 1M3D_B--1M3D_E |
| | | E | 224 | 1M3D_B--1M3D_F |
| | | | | 1M3D_C--1M3D_E |
| | | F | 223 | 1M3D_C--1M3D_F |
| | | | | 1M3D_D--1M3D_E |
| | | G | 225 | 1M3D_D--1M3D_F |
| | | | | 1M3D_D--1M3D_H |
| | | H | 224 | 1M3D_E--1M3D_F |
| | | | | 1M3D_G--1M3D_H |
| | | L | 222 | 1M3D_H--1M3D_L |
| 1M63 | 2.8 | C | 165 | 1M63_C--1M63_E |
| | | E | 359 | |
| 1MDM | 2.8 | A | 124 | 1MDM_A--1MDM_B |
| | | B | 129 | |
| 1MHW | 1.9 | A | 173 | 1MHW_A--1MHW_C |
| | | C | 41 | |
| 1NCI | 2.1 | A | 102 | 1NCI_A--1NCI_B |
| | | B | 96 | |
| 1NH2 | 1.9 | B | 46 | 1NH2_B--1NH2_C |
| | | C | 50 | |
| 1NME | 1.6 | A | 146 | 1NME_A--1NME_B |
| | | B | 92 | |
| 1NT2 | 2.9 | A | 209 | 1NT2_A--1NT2_B |
| | | B | 236 | |
| 1PON | NMR | A | 34 | 1PON_A--1PON_B |
| | | B | 34 | |
| 1PYO | 1.65 | A | 159 | 1PYO_A--1PYO_B |
| | | B | 98 | |
| 1Q68 | NMR | A | 38 | 1Q68_A--1Q68_B |
| | | B | 29 | |
| 1R4M | 3.0 | B | 418 | 1R4M_B--1R4M_I |
| | | I | 76 | |
| 1RF3 | 3.5 | A | 192 | 1RF3_A--1RF3_B |
| | | B | 24 | |
| 1RKC | 2.7 | A | 258 | 1RKC_A--1RKC_B |
| | | B | 26 | |
| 1RVF | 4.0 | 2 | 255 | 1RVF_2--1RVF_3 |
| | | 3 | 236 | 1RVF_2--1RVF_4 |
| | | 4 | 40 | 1RVF_3--1RVF_4 |
| 1SHW | 2.2 | A | 138 | 1SHW_A--1SHW_B |
| | | B | 181 | |

**Table S1.** (continued next page)

| PDB | Resolution | Chains | Nb residues | Couples |
|---|---|---|---|---|
| 1US7 | 2.3 | A<br>B | 207<br>194 | 1US7_A--1US7_B |
| 1VYH | 3.4 | A<br>B | 218<br>310 | 1VYH_A--1VYH_C |
| 1WSU | 2.3 | A<br>B<br>C<br>D | 124<br>122<br>102<br>121 | 1WSU_A--1WSU_B<br>1WSU_C--1WSU_D |
| 1Y8N | 2.6 | A<br>B | 374<br>97 | 1Y8N_A--1Y8N_B |
| 1YA5 | 2.44 | A<br>T | 198<br>89 | 1YA5_A--1YA5_T |
| 1YDI | 1.8 | A<br>B | 256<br>24 | 1YDI_A--1YDI_B |
| 1YK1 | 2.9 | A<br>E | 394<br>21 | 1YK1_A--1YK1_E |
| 1YY9 | 2.6 | C<br>D | 211<br>220 | 1YY9_C--1YY9_D |
| 1ZSG | NMR | A<br>B | 65<br>22 | 1ZSG_A--1ZSG_B |
| 1ZTP | 2.5 | A<br>C | 225<br>218 | 1ZTP_A--1ZTP_C |
| 2AGH | NMR | A<br>B<br>C | 25<br>87<br>31 | 2AGH_A--2AGH_B<br>2AGH_B--2AGH_C |
| 2AWW | 2.21 | A<br>B | 91<br>91 | 2AWW_A--2AWW_B |
| 2BDN | 2.53 | H<br>L | 217<br>214 | 2BDN_H--2BDN_L |
| 2BKI | 2.9 | B<br>D | 145<br>78 | 2BKI_B--2BKI_D |
| 2BOV | 2.66 | A<br>B | 174<br>208 | 2BOV_A--2BOV_B |
| 2C0L | 2.3 | A<br>B | 292<br>122 | 2C0L_A--2C0L_B |
| 2C35 | 2.7 | A<br>B | 129<br>171 | 2C35_A--2C35_B |
| 2C63 | 2.15 | A<br>B<br>C<br>D | 233<br>233<br>233<br>233 | 2C63_A--2C63_B<br>2C63_A--2C63_C<br>2C63_B--2C63_D<br>2C63_C--2C63_D |
| 2C74 | 2.7 | A<br>B | 235<br>234 | 2C74_A--2C74_B |
| 2C9W | 1.9 | A<br>C | 153<br>80 | 2C9W_A--2C9W_C |

**Table S1** (continued next page)

| PDB | Resolution | Chains | Nb residues | Couples |
|---|---|---|---|---|
| 2D1X | 1.9 | A<br>B<br>C | 60<br>59<br>66 | 2D1X_A--2D1X_B<br>2D1X_A--2D1X_C |
| 2DJG | 2.05 | A<br>B<br>C | 114<br>161<br>68 | 2DJG_A--2DJG_B<br>2DJG_A--2DJG_C<br>2DJG_B--2DJG_C |
| 2DRN | NMR | A<br>C | 46<br>24 | 2DRN_A--2DRN_C |
| 2DVW | 2.3 | A<br>B | 229<br>73 | 2DVW_A--2DVW_B |
| 2E9W | 3.5 | A<br>C | 468<br>132 | 2E9W_A--2E9W_C |
| 2E9X | 2.3 | A<br>B<br>C<br>D | 144<br>175<br>186<br>197 | 2E9X_A--2E9X_B<br>2E9X_A--2E9X_C<br>2E9X_A--2E9X_D<br>2E9X_B--2E9X_C<br>2E9X_B--2E9X_D<br>2E9X_C--2E9X_D |
| 2FFK | NMR | A<br>B | 242<br>69 | 2FFK_A--2FFK_B |
| 2GD4 | 3.3 | H<br>L | 234<br>54 | 2GD4_H--2GD4_L |
| 2GEZ | 2.6 | B<br>C | 133<br>166 | 2GEZ_B--2GEZ_C |
| 2GIX | 2.02 | A<br>B<br>D | 206<br>201<br>205 | 2GIX_A--2GIX_B<br>2GIX_A--2GIX_D<br>2GIX_B--2GIX_D |
| 2H0D | 2.5 | A<br>B | 97<br>100 | 2H0D_A--2H0D_B |
| 2I1N | 1.85 | A<br>B | 101<br>102 | 2I1N_A--2I1N_B |
| 2I32 | 2.7 | A<br>E | 154<br>21 | 2I32_A--2I32_E |
| 2IAE | 3.5 | A<br>B | 583<br>376 | 2IAE_A--2IAE_B |
| 2JJS | 1.85 | A<br>C | 116<br>115 | 2JJS_A--2JJS_C |
| 2JZ3 | NMR | B<br>C | 118<br>96 | 2JZ3_B--2JZ3_C |
| 2K2U | NMR | A<br>B | 115<br>35 | 2K2U_A--2K2U_B |
| 2NL9 | 1.55 | A<br>B | 140<br>23 | 2NL9_A--2NL9_B |

**Table S1** (continued next page)

| PDB | Resolution | Chains | Nb residues | Couples |
|---|---|---|---|---|
| 2NNA | 2.1 | A | 182 | 2NNA_A--2NNA_B |
| | | B | 182 | |
| 2NNW | 2.7 | A | 350 | 2NNW_A--2NNW_B |
| | | B | 227 | |
| 2NQB | 2.3 | D | 95 | 2NQB_D--2NQB_G |
| | | G | 105 | |
| 2NVU | 2.8 | B | 789 | 2NVU_B--2NVU_C |
| | | C | 176 | |
| 2O8A | 2.61 | A | 295 | 2O8A_A--2O8A_I |
| | | I | 59 | |
| 2ODB | 2.4 | A | 177 | 2ODB_A--2ODB_B |
| | | B | 35 | |
| 2OT3 | 2.1 | A | 253 | 2OT3_A--2OT3_B |
| | | B | 157 | |
| 2P1L | 2.5 | A | 141 | 2P1L_A--2P1L_B |
| | | B | 24 | |
| 2P1M | 1.8 | A | 90 | 2P1M_A--2P1M_B |
| | | B | 567 | |
| 2PAV | 1.8 | A | 361 | 2PAV_A--2PAV_P |
| | | P | 139 | |
| 2PJY | 3.0 | A | 112 | 2PJY_A--2PJY_B |
| | | B | 108 | 2PJY_A--2PJY_C |
| | | C | 79 | 2PJY_B--2PJY_C |
| 2Q7N | 4.0 | A | 480 | 2Q7N_A--2Q7N_B |
| | | B | 180 | |
| 2QFA | 1.4 | A | 137 | 2QFA_A--2QFA_B |
| | | B | 62 | 2QFA_A--2QFA_C |
| | | C | 45 | 2QFA_B--2QFA_C |
| 2QOU | 3.93 | C | 206 | 2QOU_C--2QOU_E |
| | | D | 205 | 2QOU_C--2QOU_J |
| | | E | 150 | 2QOU_C--2QOU_N |
| | | F | 100 | 2QOU_D--2QOU_E |
| | | G | 150 | 2QOU_E--2QOU_H |
| | | H | 29 | 2QOU_F--2QOU_R |
| | | I | 127 | 2QOU_G--2QOU_I |
| | | J | 98 | 2QOU_G--2QOU_K |
| | | K | 11 | 2QOU_H--2QOU_L |
| | | L | 123 | 2QOU_H--2QOU_Q |
| | | M | 114 | 2QOU_I--2QOU_J |
| | | N | 96 | 2QOU_I--2QOU_N |
| | | O | 88 | 2QOU_J--2QOU_N |
| | | Q | 80 | 2QOU_K--2QOU_R |
| | | R | 55 | 2QOU_K--2QOU_U |
| | | S | 79 | 2QOU_L--2QOU_Q |
| | | U | 51 | 2QOU_M--2QOU_S |
| | | | | 2QOU_N--2QOU_S |
| | | | | 2QOU_O--2QOU_Q |
| | | | | 2QOU_R--2QOU_U |

**Table S1** (continued next page)

| PDB | Resolution | Chains | Nb residues | Couples |
|---|---|---|---|---|
| 2QOV | 3.93 | 1 | 50 | |
| | | 3 | 64 | |
| | | 4 | 38 | 2QOV_1--2QOV_3 |
| | | D | 209 | 2QOV_3--2QOV_L |
| | | E | 201 | 2QOV_4--2QOV_G |
| | | G | 176 | 2QOV_D--2QOV_K |
| | | H | 149 | 2QOV_D--2QOV_N |
| | | K | 121 | 2QOV_D--2QOV_P |
| | | L | 143 | 2QOV_E--2QOV_L |
| | | M | 136 | 2QOV_G--2QOV_H |
| | | N | 120 | 2QOV_H--2QOV_Z |
| | | P | 114 | 2QOV_K--2QOV_P |
| | | T | 93 | 2QOV_M--2QOV_V |
| | | V | 94 | 2QOV_T--2QOV_X |
| | | X | 63 | |
| | | Z | 77 | |
| 2R9P | 1.4 | A | 24 | 2R9P_A--2R9P_E |
| | | E | 58 | |
| 2RGN | 3.5 | A | 324 | 2RGN_A--2RGN_B |
| | | B | 327 | 2RGN_B--2RGN_C |
| | | C | 177 | |
| 2RHK | 1.95 | A | 119 | 2RHK_A--2RHK_C |
| | | C | 63 | |
| 2RMK | NMR | A | 192 | 2RMK_A--2RMK_B |
| | | B | 81 | |
| 2UZI | 2.0 | H | 114 | 2UZI_H--2UZI_L |
| | | L | 104 | |
| 2V17 | 1.65 | H | 222 | 2V17_H--2V17_L |
| | | L | 214 | |
| 2V8Q | 2.1 | A | 102 | 2V8Q_A--2V8Q_B |
| | | B | 73 | 2V8Q_A--2V8Q_E |
| | | E | 304 | 2V8Q_B--2V8Q_E |
| 2VGL | 2.59 | A | 600 | 2VGL_A--2VGL_B |
| | | | | 2VGL_A--2VGL_M |
| | | B | 579 | 2VGL_A--2VGL_S |
| | | M | 396 | 2VGL_B--2VGL_M |
| | | | | 2VGL_B--2VGL_S |
| | | S | 142 | 2VGL_M--2VGL_S |
| 2VP7 | 1.65 | A | 66 | 2VP7_A--2VP7_B |
| | | B | 33 | |
| 2Z3Q | 1.85 | B | 81 | 2Z3Q_B--2Z3Q_C |
| | | C | 117 | |
| 2Z5H | 2.89 | B | 51 | 2Z5H_B--2Z5H_I |
| | | I | 39 | 2Z5H_I--2Z5H_T |
| | | T | 34 | |

**Table S1** (continued next page)

| PDB | Resolution | Chains | Nb residues | Couples |
|---|---|---|---|---|
| 2ZCH | 2.83 | H | 229 | 2ZCH_H--2ZCH_L |
| | | L | 215 | 2ZCH_H--2ZCH_P |
| | | P | 237 | 2ZCH_L--2ZCH_P |
| 2ZL1 | 2.0 | A | 119 | 2ZL1_A--2ZL1_B |
| | | B | 116 | |
| 3B6F | 3.45 | C | 106 | 3B6F_C--3B6F_H |
| | | H | 99 | |
| 3BC1 | 1.8 | B | 52 | 3BC1_B--3BC1_E |
| | | E | 175 | |
| 3BES | 2.2 | L | 133 | 3BES_L--3BES_R |
| | | R | 250 | |
| 3BJ4 | 2.0 | A | 37 | 3BJ4_A--3BJ4_B |
| | | B | 38 | |
| 3BPL | 2.93 | B | 202 | 3BPL_B--3BPL_C |
| | | C | 194 | |
| 3BRT | 2.25 | B | 61 | 3BRT_B--3BRT_C |
| | | C | 43 | |
| 3BRW | 3.4 | B | 337 | 3BRW_B--3BRW_D |
| | | D | 167 | |
| 3BS5 | 2.0 | A | 83 | 3BS5_A--3BS5_B |
| | | B | 74 | |
| 3BT2 | 2.5 | A | 124 | 3BT2_A--3BT2_U |
| | | B | 40 | 3BT2_B--3BT2_U |
| | | H | 212 | 3BT2_H--3BT2_L |
| | | L | 211 | 3BT2_H--3BT2_U |
| | | U | 259 | 3BT2_L--3BT2_U |
| 3BYH | 12.0 (EM) | A | 374 | 3BYH_A--3BYH_B |
| | | B | 231 | |
| 3C08 | 2.15 | H | 217 | 3C08_H--3C08_L |
| | | L | 206 | |
| 3C5J | 1.8 | A | 178 | 3C5J_A--3C5J_B |
| | | B | 182 | |
| 3C66 | 2.6 | A | 529 | 3C66_A--3C66_C |
| | | C | 24 | |
| 3CH5 | 2.1 | A | 193 | 3CH5_A--3CH5_B |
| | | B | 37 | |
| 3CL3 | 3.2 | A | 172 | 3CL3_A--3CL3_D |
| | | D | 59 | |

**Table S1** (continued next page)

| PDB | Resolution | Chains | Nb residues | Couples |
|---|---|---|---|---|
| 3CWB | 3.51 | A | 443 | 3CWB_A--3CWB_B |
| | | B | 421 | 3CWB_A--3CWB_D |
| | | D | 241 | 3CWB_A--3CWB_E |
| | | E | 196 | 3CWB_A--3CWB_J |
| | | F | 100 | 3CWB_D--3CWB_E |
| | | H | 70 | 3CWB_D--3CWB_F |
| | | J | 61 | 3CWB_D--3CWB_H |
| | | P | 379 | 3CWB_D--3CWB_J |
| | | T | 79 | 3CWB_E--3CWB_J |
| | | | | 3CWB_E--3CWB_P |
| | | | | 3CWB_P--3CWB_T |
| 3CXE | 3.3 | A | 412 | 3CXE_A--3CXE_B |
| | | B | 105 | 3CXE_B--3CXE_C |
| | | C | 116 | |
| 3D0G | 2.8 | A | 597 | 3D0G_A--3D0G_E |
| | | E | 173 | |
| 3D1M | 1.7 | A | 148 | 3D1M_A--3D1M_D |
| | | D | 99 | |
| 3D2U | 2.21 | A | 281 | 3D2U_A--3D2U_B |
| | | B | 99 | |
| 3D48 | 2.5 | P | 165 | 3D48_P--3D48_R |
| | | R | 195 | |
| 3D85 | 1.9 | A | 213 | 3D85_A--3D85_B |
| | | B | 216 | 3D85_A--3D85_C |
| | | C | 133 | 3D85_B--3D85_C |
| | | D | 290 | 3D85_C--3D85_D |
| 3DGC | 2.5 | M | 141 | 3DGC_S--3DGC_M |
| | | S | 207 | |

**Table S1.** List of PDB structures used to create the CC-D dataset with its corresponding resolution. The chains included in the dataset, the number of residues of each chain and the experimental partners couples are indicated in the third, fourth and fifth columns respectively.

| protein | PrimI | OR (%) | SecI | OR (%) | PrimI/SecI |
|---|---|---|---|---|---|
| 1AIK_C | 1AIK_C--1AIK_A | 69 | 1AIK_C--1AIK_N | 31 | 2.2 |
| 1AOX_A | 1AOX_A--homolog1V7PB1 | 59 | 1AOX_A--1AOX_B | 20 | 2.95 |
| 1AOX_B | 1AOX_B--homolog1V7PB2 | 68 | 1AOX_B--1AOX_A | 7 | 9.71 |
| 1EF1_C | 1EF1_C--1EF1_A | 89 | 1EF1_C--1EF1_D | 10 | 8.90 |
| 1FNT_J | 1FNT_J--1FNT_I | 33 | 1FNT_J--1FNT_B | 8 | 4.13 |
| 1GC1_C | 1GC1_C--homolog1CDH | 93 | 1GC1_C--1GC1_G | 7 | 13.29 |
| 1GK4_A | 1GK4_A--1GK4_B | 77 | 1GK4_A--1GK4_F | 12 | 6.42 |
| 1GK4_C | 1GK4_C--1GK4_D | 85 | 1GK4_C--1GK4_F | 13 | 6.54 |
| 1GK4_D | 1GK4_D--1GK4_C | 84 | 1GK4_D--1GK4_F | 14 | 6.00 |
| 1GK4_F | 1GK4_F--1GK4_E | 66 | 1GK4_F--1GK4_D | 22 | 3.00 |
| 1H2K_A | 1H2K_A--1H2K_B | 73 | 1H2K_A--1H2K_S | 27 | 2.70 |
| 1I7X_A | 1I7X_A--homolog1G3J | 58 | 1I7X_A--1I7X_C | 19 | 3.05 |
| 1JJO_A | 1JJO_A--1JJO_C | 70 | 1JJO_A--1JJO_E | 22 | 3.18 |
| 1JJO_E | 1JJO_E--1JJO_C | 62 | 1JJO_E--1JJO_A | 38 | 1.63 |
| 1KFU_L | 1KFU_L--homolog3DF0 | 96 | 1KFU_L--1KFU_S | 4 | 24.00 |
| 1LDK_A | 1LDK_A--1LDK_E | 64 | 1LDK_A--1LDK_B | 11 | 5.82 |
| 1LDK_B | 1LDK_B--1LDK_C | 78 | 1LDK_B--1LDK_A | 22 | 3.55 |
| 1LI1_A | 1LI1_A--1LIA_C | 73 | 1LI1_A--1LI1_B | 22 | 3.32 |
| 1LI1_B | 1LI1_B--1LI1_A | 63 | 1LI1_B--1LI1_C | 30 | 2.10 |
| 1M3D_A | 1M3D_A--1M3D_C | 63 | 1M3D_A--1M3D_B | 33 | 1.91 |
| 1M3D_B | 1M3D_B--1M3D_A | 55 | 1M3D_B--1M3D_F | 1 | 55.00 |
| 1M3D_C | 1M3D_C--1M3D_B | 49 | 1M3D_C--1M3D_E | 1 | 49.00 |
| 1M3D_E | 1M3D_E--1M3D_D | 57 | 1M3D_E--1M3D_C | 1 | 57.00 |
| 1M3D_F | 1M3D_F--1M3D_D | 44 | 1M3D_F--1M3D_B | 1 | 44.00 |
| 1M3D_G | 1M3D_G--1M3D_I | 60 | 1M3D_G--1M3D_H | 35 | 1.71 |
| 1M3D_L | 1M3D_L--1M3D_J | 44 | 1M3D_L--1M3D_H | 1 | 44.00 |
| 1NH2_B | 1NH2_B--1NH2_D | 81 | 1NH2_B--1NH2_C | 19 | 4.26 |
| 1NH2_C | 1NH2_C--1NH2_D | 44 | 1NH2_C—1NH2_B | 16 | 2.75 |
| 1PYO_B | 1PYO_B--1PYO_A | 88 | 1PYO_B--1PYO_D | 6 | 14.67 |
| 1R4M_B | 1R4M_B--1R4M_A | 60 | 1R4M_B--1R4M_I | 20 | 3.00 |
| 1R4M_I | 1R4M_I--1R4M_B | 100 | 1R4M_I--1R4M_A | 0 | NA |
| 1RF3_A | 1RF3_A--1RF3_E | 44 | 1RF3_A--1RF3_B | 26 | 1.69 |
| 1VYH_A | 1VYH_A--1VYH_B | 72 | 1VYH_A--1VYH_C | 13 | 5.54 |
| 1YY9_C | 1YY9_C--1YY9_D | 98 | 1YY9_C--1YY9_A | 2 | 49.00 |
| 1YY9_D | 1YY9_D--1YY9_C | 62 | 1YY9_D--1YY9_A | 38 | 1.63 |
| 2BDN_H | 2BDN_H--2BDN_L | 64 | 2BDN_H--2BDN_A | 36 | 1.78 |
| 2BDN_L | 2BDN_L--2BDN_H | 92 | 2BDN_L—2BDN_A | 8 | 11.5 |
| 2BKI_B | 2BKI_B--2BKI_A | 63 | 2BKI_B--2BKI_D | 37 | 1.70 |
| 2BKI_D | 2BKI_D--2BKI_B | 63 | 2BKI_D--2BKI_A | 37 | 1.70 |
| 2BOV_A | 2BOV_A--homolg1UAD | 79 | 2BOV_A--2BOV_B | 21 | 3.76 |
| 2C0L_A | 2C0L_A--homolog1FCH | 80 | 2C0L_A--2C0L_B | 20 | 4.00 |
| 2C74_A | 2C74_A--2C74_P | 81 | 2C74_A--2C74_B | 19 | 4.26 |
| 2C74_B | 2C74_B--2C74_Q | 74 | 2C74_B--2C74_A | 26 | 2.85 |
| 2C9W_C | 2C9W_C--2C9W_B | 77 | 2C9W_C--2C9W_A | 23 | 3.35 |

**Table S2** (continued next page)

| protein | PrimI | OR (%) | SecI | OR (%) | PrimI/SecI |
|---|---|---|---|---|---|
| 2DJG_A | 2DJG_A--2DJG_B | 78 | 2DJG_A--2DJG_C | 22 | 3.55 |
| 2DJG_B | 2DJG_B--2DJG_A | 71 | 2DJG_B--2DJG_C | 29 | 2.45 |
| 2DRN_A | 2DRN_A--2DRN_B | 72 | 2DRN_A--2DRN_C | 28 | 2.57 |
| 2E9W_A | 2E9W_A--2E9W_B | 92 | 2E9W_A--2E9W_C | 8 | 11.50 |
| 2E9X_A | 2E9X_A--2E9X_D | 62 | 2E9X_A--2E9X_F | 9 | 6.89 |
| 2E9X_C | 2E9X_C--2E9X_A | 58 | 2E9X_C--2E9X_B | 16 | 3.63 |
| 2GEZ_B | 2GEZ_B--2GEZ_A | 71 | 2GEZ_B--2GEZ_C | 13 | 5.46 |
| 2GEZ_C | 2GEZ_C--2GEZ_D | 46 | 2GEZ_C--2GEZ_B | 5 | 9.20 |
| 2NNW_A | 2NNW_A--homolog3NMU | 84 | 2NNW_A--2NNW_B | 5 | 16.80 |
| 2NQB_G | 2NQB_G--2NQB_H | 71 | 2NQB_G--2NQB_D | 9 | 7.89 |
| 2O8A_I | 2O8A_I--homolog3N5U | 83 | 2O8A_I--2O8A_A | 17 | 4.88 |
| 2P1L_B | 2P1L_B--homologH | 64 | 2P1L_B--2P1L_A | 36 | 1.78 |
| 2PJY_C | 2PJY_C--2PJY_D | 57 | 2PJY_C--2PJY_A | 22 | 2.59 |
| 2RGN_B | 2RGN_B--2RGN_C | 68 | 2RGN_B--2RGN_A | 32 | 2.13 |
| 2RHK_A | 2RHK_A--2RHK_B | 73 | 2RHK_A--2RHK_C | 27 | 2.70 |
| 2UZI_H | 2UZI_H--2UZI_L | 84 | 2UZI_H--2UZI_R | 16 | 5.25 |
| 2UZI_L | 2UZI_L--2UZI_H | 75 | 2UZI_L--2UZI_R | 25 | 3.00 |
| 2V8Q_A | 2V8Q_A--2V8Q_B | 80 | 2V8Q_A--2V8Q_E | 20 | 4.00 |
| 2V8Q_E | 2V8Q_E--2V8Q_A | 74 | 2V8Q_E--2V8Q_B | 26 | 2.85 |
| 2VGL_B | 2VGL_B--2VGL_M | 58 | 2VGL_B--2VGL_S | 7 | 8.29 |
| 2VGL_M | 2VGL_M--2VGL_B | 76 | 2VGL_M--2VGL_S | 16 | 4.75 |
| 2VGL_S | 2VGL_S--2VGL_A | 67 | 2VGL_S—2VGL_B | 18 | 3.72 |
| 2Z3Q_C | 2Z3Q_C--2Z3Q_D | 62 | 2Z3Q_C--2Z3Q_B | 38 | 1.63 |
| 2Z5H_B | 2Z5H_B--2Z5H_A | 48 | 2Z5H_B--2Z5H_I | 13 | 3.69 |
| 2ZCH_H | 2ZCH_H--2ZCH_L | 83 | 2ZCH_H--2ZCH_P | 17 | 4.88 |
| 2ZCH_L | 2ZCH_L--2ZCH_H | 89 | 2ZCH_L--2ZCH_P | 11 | 8.09 |
| 3B6F_C | 3B6F_C--3B6F_D | 68 | 3B6F_C--3B6F_E | 12 | 5.67 |
| 3BC1_E | 3BC1_E--3BC1_F | 64 | 3BC1_E--3BC1_B | 9 | 7.11 |
| 3BES_L | 3BES_L--3BES_A | 56 | 3BES_L--3BES_R | 27 | 2.07 |
| 3BES_R | 3BES_R--3BES_D | 31 | 3BES_R—3BES_L | 7 | 4.43 |
| 3BT2_H | 3BT2_H--3BT2_U | 68 | 3BT2_H--3BT2_L | 32 | 2.13 |
| 3BT2_L | 3BT2_L--3BT2_H | 70 | 3BT2_L--3BT2_U | 30 | 2.33 |
| 3CH5_A | 3CH5_A—homolog_1IBR | 73 | 3CH5_A--3CH5_B | 27 | 2.70 |
| 3CL3_D | 3CL3_D—3CL3_B | 68 | 3CL3_D--3CL3_A | 13 | 5.23 |
| 3CWB_B | 3CWB_B--3CWB_O | 86 | 3CWB_B--3CWB_A | 3 | 28.67 |
| 3CWB_E | 3CWB_E--3CWB_P | 66 | 3CWB_E--3CWB_D | 7 | 9.43 |
| 3CWB_F | 3CWB_F--3CWB_C | 68 | 3CWB_F--3CWB_D | 15 | 4.53 |
| 3D85_A | 3D85_A--3D85_B | 96 | 3D85_A--3D85_C | 4 | 24.00 |
| 3D85_B | 3D85_B--3D85_A | 67 | 3D85_B--3D85_C | 33 | 2.03 |
| 3D85_C | 3D85_C--3D85_D | 58 | 3D85_C--3D85_B | 10 | 5.80 |
| 3DGC_M | 3DGC_M--3DGC_S | 70 | 3DGC_M--3DGC_R | 30 | 2.33 |

**Table S2.** List of proteins presenting occupancy rates (OR) at least 50% superior for the Primary Interface (PrimI) compared to Secondary Interface (SecI). The last column PrimI/SecI indicates the ratio value of the PrimI's occupancy rate over the SecI's occupancy rate.

| Descriptors | Greater | Less |
|---|---|---|
| % Acidic | 7.21E-01 | 2.79E-01 |
| % Acyclic | 3.43E-01 | 6.57E-01 |
| % Aliphatic | 1.44E-02 | 9.86E-01 |
| % Alpha character | 2.55E-01 | 7.45E-01 |
| % Aromatic | 6.97E-01 | 3.03E-01 |
| % Basic | 8.89E-01 | 1.11E-01 |
| % Beta character | 1.38E-02 | 9.86E-01 |
| Carbon | 1.24E-02 | 9.88E-01 |
| % Charged contribution | 9.81E-01 | 1.95E-02 |
| % Charged Residues | 8.55E-01 | 1.45E-01 |
| Circularity | 2.70E-01 | 7.30E-01 |
| % Coil | 8.34E-01 | 1.66E-01 |
| ContRes | **1.09E-04** | 1.00E+00 |
| ContRC | **1.61E-08** | 1.00E+00 |
| ContRHyd | **8.74E-10** | 1.00E+00 |
| ContRN | **2.54E**-08 | 1.00E+00 |
| ContRP | **8.18E-06** | 1.00E+00 |
| % Cyclic | 6.57E-01 | 3.43E-01 |
| Eccentricity | 7.87E-01 | 2.13E-01 |
| Fluor | NA | NA |
| Gap Volume | **6.30E-05** | 1.00E+00 |
| Hydrogen | 5.91E-01 | 4.09E-01 |
| % Interface Accessible Surface Area | **1.71E-09** | 1.00E+00 |
| Interface Accessible Surface Area | **1.03E-08** | 1.00E+00 |
| % Large | 4.23E-01 | 5.77E-01 |
| % Medium | 2.29E-01 | 7.71E-01 |
| Nb of hydrogen bonds | **1.09E-03** | 9.99E-01 |
| Nb of non-bonded contacts | **1.55E-06** | 1.00E+00 |
| Nb of salt bridges | 1.61E-01 | 8.39E-01 |
| % Neutral contribution | 9.19E-02 | 9.08E-01 |
| (N)itrogen | 9.37E-01 | 6.30E-02 |
| % Non polar contribution | **2.60E-08** | 1.00E+00 |
| N+O+P+S | 3.13E-01 | 6.87E-01 |
| Number of Segments | **8.66E-05** | 1.00E+00 |
| (O)xygen | 9.26E-01 | 7.44E-02 |
| (P)hosphorus | NA | NA |
| Planarity | **1.61E-07** | 1.00E+00 |
| % Polar contribution | 1.00E+00 | **2.61E-08** |
| % Small | 8.96E-01 | 1.04E-01 |
| (S)ulfur | 1.86E-01 | 8.14E-01 |
| Total Interface Area | **1.88E-08** | 1.00E+00 |
| Total Nb of Disulfide bonds | 7.91E-01 | 2.09E-01 |
| Total Nb of Segments | **3.25E-06** | 1.00E+00 |

**Table S3**. p-values obtained for each one of the 43 2P2I inspector descriptors in the one-tailed Student test with the option « greater » (column « Greater ») and « less » (column « Less »). The significant p-values using the Bonferonni threshold are indicated in bold.
